# Nanoscale cluster mapping of molecular complexes with pixel-based correlations

**DOI:** 10.1101/2024.07.02.601646

**Authors:** Tai Kiuchi, Ryouhei Kobayashi, Shuichiro Ogawa, Louis L. H. Elverston, Naoki Watanabe

**Author notes:** Correspondence should be addressed to T.K. Nagahama Institute of Bio-Science and Technology, 1266 Tamura-cho, Nagahama, Shiga 526-0829, Japan (Shuichiro Ogawa).

## Abstract

Super-resolution microscopy achieves a few nanometers resolution, but protein quantification and colocalization is limited by its labeling density. Here we present a method to map molecular complexes using the multiplexed data of IRIS super-resolution imaging. We developed antiserum-derived Fab IRIS probes for high-density labeling of endogenous targets, and image analysis, Protein Cluster coloring (PC-coloring), which employs pixel-based principal component analysis and clustering. PC-coloring was first used to map regions of distinct ratios of targets on the image. The molecular complex formation was then evaluated by correlation between all combinations of two proteins in each PC-colored region. We elucidate receptor recruitment and the complex formation in a clathrin-coated structure (CCS) which is pre-assembled prior to endocytosis. Upon EGF stimulation, EGFR-, EGFR-Grb2-complex- and Grb2-dominant regions lined up from the CCS rim. Along the interior of Grb2-dominant regions, Eps15, FCHo1/2 and intersectin-1 formed a complex with Grb2. The interior of Eps15-FCHo1/2 complexes was lined with Eps15-intersectin-1 complexes. The results reveal a laminar complex formation of EGFR, Grb2 and CCS components at the recruitment sites.

---

Super-resolution microscopy achieves a resolution of a few nanometers, allowing visualization of molecular localization within the complex^1,2^. However, multiple labeling of endogenous proteins with antibodies is impossible in a region smaller than the antibody size (> 10 nm)^3,4^. Especially in multiplexed super-resolution microscopy, physical interference between antibodies may reduce the accessibility to the antigens as previously reported^5^. Due to the incomplete labeling, accuracy in colocalization analysis cannot be improved solely by increasing the resolution. To overcome the labeling problem, we developed a multiplexed super-resolution microscopy, named image reconstruction by integrating exchangeable single-molecule localization (IRIS)^6^. IRIS employs fluorescent probes (IRIS probes) that rapidly associate with and dissociate from the endogenous target^6^. The fast exchange property enables high-density and sequential labeling of multiple targets without causing physical interference between the probes^5,6^. The IRIS probes have ever been produced from protein fragments, Fab fragments of mouse monoclonal antibodies, and engineered antibody and nanobody fragments^5–7^. Here we introduce a new IRIS probe from antiserum. The antiserum contains polyclonal antibodies against multiple epitopes in the target protein. Therefore, antiserum-derived Fab probes are expected to stain targets, some epitopes of which are not exposed in a complex. Furthermore, to map molecular complexes between multiple targets, we present Protein Cluster coloring (PC-coloring) that employs principal component analysis (PCA) and clustering of protein-size pixels in IRIS images.

In the canonical model of clathrin-mediated endocytosis (CME), clathrin-coated pits (CCPs) are formed, grow to 150 – 200 nm in diameter upon binding to the receptor cargoes, and bud off^8^. However, electron microscopy studies show clathrin-coated structures (CCSs) ranging in size from tens nanometers to a few micrometers, including CCPs and clathrin plaque^9–13^. CCPs are not isolated structures but are often linked to the rim of a clathrin plaque^10,11,13^. Clathrin plaques also often contain curved clathrin lattices like a CCP^10^. Fluorescence microscopy studies show hotspots of CME and EGFR recruitment upon EGF stimulation in CCSs^13,14^. Hence, CCS is a molecular apparatus which is pre-assembled for receptor recruitment prior to CME. Although the interaction network between the CCS components and the recruited receptors have been proposed^15^, their spatial relationship is poorly understood. Many immunoelectron microscopy and super-resolution microscopy studies have observed the localization of a single target and have shown its averaged distribution in many CCSs^16–20^. However, due to the heterogeneity of the CCSs, colocalization between multiple molecules (i.e. molecular complexes) remains to be determined.

## Spatial relationship among CCS-associated proteins

We visualized the distribution of eight endogenous CCS-associated proteins, which were clathrin light chain (CLC), adaptor protein 2 complex beta subunit (AP2β2), EGFR, Grb2, transferrin receptors (TfR), FCHo1 and 2 (FCHo1/2), Eps15 and intersectin-1 (ITSN1). These proteins were localized in CCSs visualized by CLC and AP2β2 (Fig. 1A). We produced the Fab probes from antiserum and verified their labeling to the endogenous targets in HeLa cells (Supplementary Figs. 1–8, Supplementary Table 1, and Methods). The average labeling densities of these probes were 2.9 – 4.8 labels per 10 nm pixel after correcting for overcounting of probe binding across multiple frames (Extended Data Fig. 1D). These densities exceed the maximum occupied density (0.9 per 10 nm pixel) of an antibody (12 nm in size) or an EGFR dimer (11.9 nm in size^21^). We further improved the localization accuracy using normalized Gaussian with mean localization and standard error (SE) of a probe bound over multiple frames^2^ (Methods). In the localization distribution of a long-binding probe, the normalized Gaussian represents the probability distribution of the true target location, according to the central limit theorem (Fig. 1B, bottom graph). The SE decreased with the number of the localizations, reaching around 1 nm (Extended Data Fig. 2A). The mode of the SEs was reduced to 10.7 nm compared to 24.2 nm in the standard deviations (SDs) of the localization distribution (Extended Data Fig. 2B, C). The resolution of the Gaussian rendering IRIS image with mean localization is 2.26 times better than that of the IRIS image reconstructed by integrating the localizations (Fig. 1C and Extended Data Fig. 3). In the Gaussian rendering image, the labeling densities in the target-localized area were 1.5 – 6.6 or higher labels per 10 nm pixel (Extended Data Fig. 1E).

**Fig. 1.**
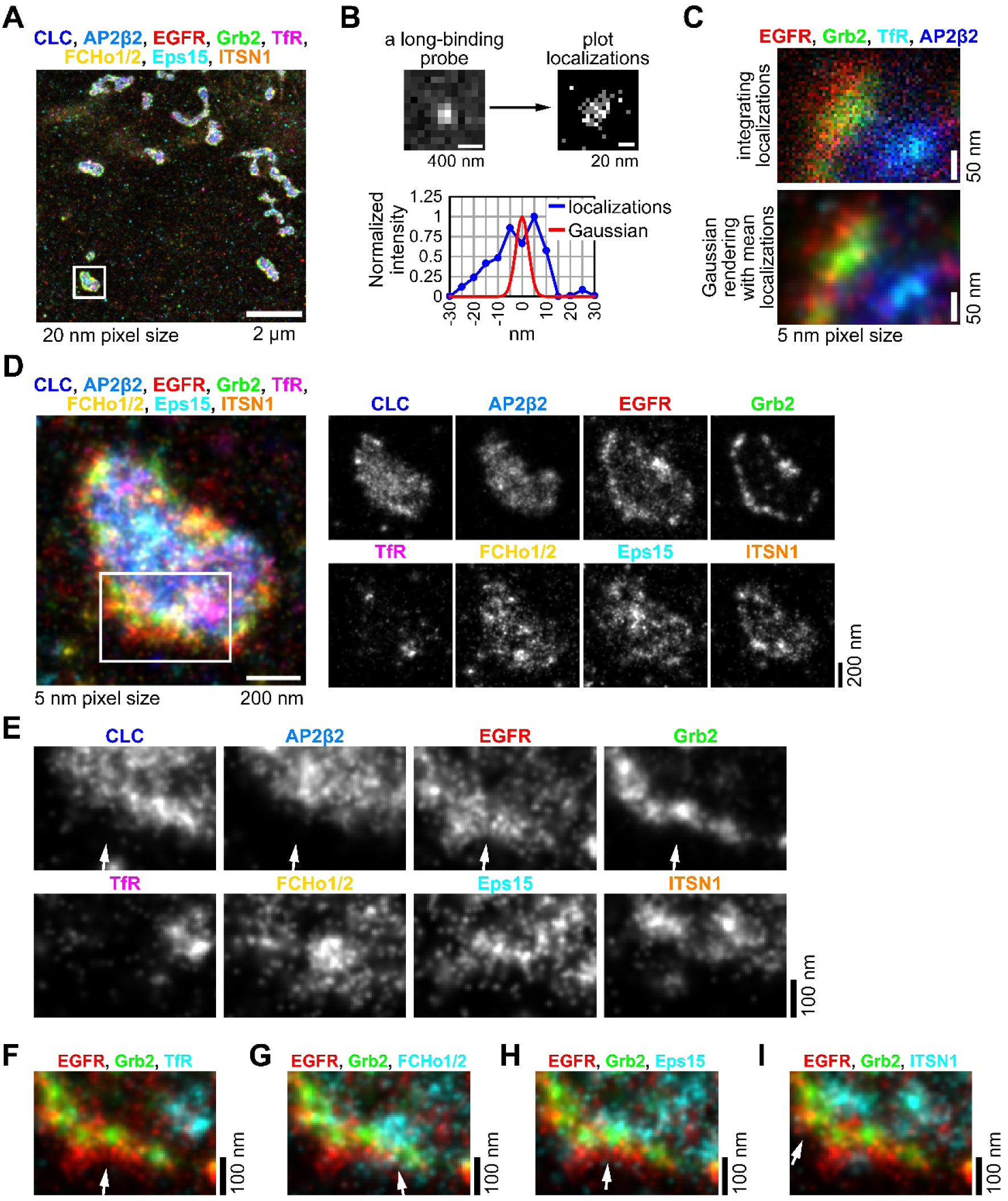
Multiplexed IRIS super-resolution image of eight endogenous targets. (**A**) The merged IRIS image of eight targets reconstructed by integrating localizations at 20 nm sized pixel. HeLa cells were stimulated with 100 ng/ml EGF for 20 min at room temperature. (**B**) Improved localization accuracy using normalized Gaussian with mean localization. A fluorescent speckle of a long-binding probe (upper left). IRIS image reconstructed by integrating localizations of the long-binding probe at 5 nm sized pixel (upper right). In the bottom graph, the cross-sectional profile of the reconstructed image (blue) was plotted to show the distribution of the localizations, whose standard deviation indicates the localization accuracy. A normalized two-dimensional Gaussian with mean and SE of the localizations (red) were plotted to show the accuracy of the mean localization. (**C**) Merged IRIS images reconstructed by integrating localizations (upper) and by Gaussian rendering with mean localizations (lower) in the part of the boxed area shown in **A**. (**D**) Spatial relationship of the eight targets in the CCS in the box area shown in **A**. The IRIS image was reconstructed by Gaussian rendering with mean localizations at 5 nm sized pixel. (**E**) The enlarged image of the box area shown in **D**. (**F**–**I**) The merged IRIS image of three targets in **E**. The target names are indicated in the panel.

To investigate molecular complexes at EGFR recruitment sites in CCSs, cells were stimulated with a saturated concentration (100 ng/ml) of EGF at room temperature for delayed endocytosis^22^. Under this condition, EGFR-tagRFP was recruited in the pre-assembled CCSs (Supplementary Fig. S2A, B). The IRIS images with improved resolution revealed clusters of the eight proteins in a single CCS (Fig. 1D). EGFR and Grb2 clusters were localized at the outer rim of the CCS, the width of which ranged from around 50 nm to 100 nm (Fig. 1E, F, arrows). FCHo1/2, Eps15 and ITSN1 clusters were found at the interior of Grb2 clusters (Fig. 1G–I, arrows). Interestingly, EGFR clusters were more extended outward at the CCS rim than Grb2 clusters (Fig. 1F, arrow), suggesting either stepwise formation of EGFR-Grb2 complex or the recruitment of Grb2 by a molecule(s) other than EGFR.

Before EGF stimulation, EGFR was spread across the plasma membrane, and Grb2 was not localized in the CCSs (Extended Data Fig. 4A). After 3 min of EGF simulation at 37℃, EGFR and Grb2 clusters were localized at CCS rims (Extended Data Fig. 4B, yellow arrowheads). However, at 5 min, the number of EGFR clusters decreased in many CCSs (Extended Data Fig. 4C, cyan arrowheads), suggesting internalization of EGFR. Interestingly many Grb2 clusters remained at the CCS rims, suggesting Grb2 complexes without EGFR (Extended Data Fig. 4C, green arrowheads). These clusters also colocalized with mCherry (mCh)-SOS1 (Ras GEF) and cCbl-mCh (E3 ligase of EGFR^23^) in IRIS images with anti-mCh nanobody^24^ (Extended Data Fig. 5C, D). These results suggest that EGFR and Grb2 clusters may contain molecular complexes of different compositions at the EGFR recruitment sites.

## PC-coloring for molecular complexes

To map the molecular complexes at the EGFR recruitment sites, we developed an image analysis, Protein Cluster coloring (PC-coloring). PC-coloring classifies pixels in the merged IRIS image by PCA and clustering based on the ratio of the standardized (z-score) intensities of multiple targets (Fig. 2A, top panels). A PCA score image was created by plotting the raw intensity of the targets (Fig. 2A, top left panel) in the coordinates of the first and second principal components (PCs) (Fig. 2a, top center panel). In the PCA score image, EGFR (red)-dominant, Grb2 (green)-dominant, Eps15 (cyan)-dominant and AP2β2 (blue)-dominant pixels were located away from the center of the image. The pixels were then clustered using k-means method, a commonly used clustering method (Fig. 2A, top right panel). The clustered pixels are shown by eight-colored circles on a PCA cluster image (Fig. 2A, bottom left panel). This coloring was performed based on the similarity of the target intensity ratio in the dendrogram, the mean z-score intensities of the target (Fig. 2A, middle panels) and the position of the clustered pixels in the PCA cluster image (Fig. 2A, bottom left panel and Methods). The colors of the clustered pixels were then assigned on the original IRIS image, generating a super-resolved (SR) cluster image (Fig. 2A, bottom center panel). In the SR cluster image, EGFR-dominant (red), EGFR- and Grb2-dominant (yellow), Grb2-dominant (green) pixels and orange pixels were localized in this order from the outside of the CCS rim (Fig. 2B, C).

**Fig. 2.**
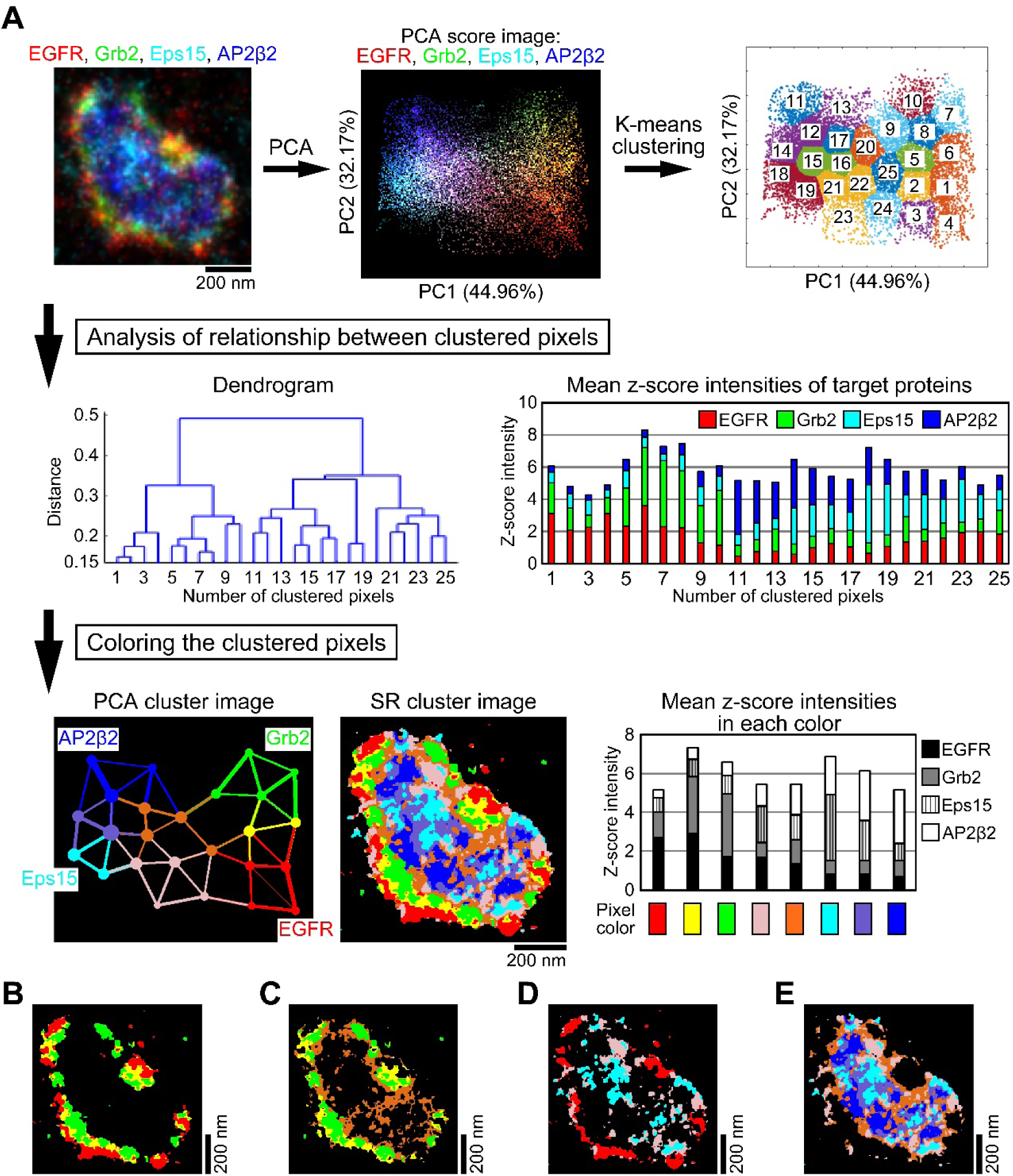
Colocalization analysis by PC-coloring. (**A**) Outline of PC-coloring (also see Methods). The merged IRIS image of EGFR (red), Grb2 (green), Eps15 (cyan) and AP2β2 (blue) shown in Fig. 1D (top left). The pixels in the IRIS image are classified by principal component analysis (PCA) based on the ratios of the standardized (z-score) intensities. A PCA score image is created by plotting the raw intensity of the target of the IRIS image (top left) in the coordinates of the first and second principal components (PC1, PC2) (top center). These pixels are clustered by k-means method (top right). A dendrogram is created based on the similarity of the intensity ratio in the clustered pixels (middle left). The mean z-score intensity of each target is calculated in the clustered pixels (middle right). The number of the clustered pixels in the top right panel is the same as the number shown in the dendrogram. The clustered pixels are shown by eight-colored circles in the coordinates of the first and second PCs as a PCA cluster image (bottom left). The center of the circle indicates that of the clustered pixels. Its area is proportional to the number of pixels in the clustered pixels. These circles are connected by lines indicating the proximity between the clustered pixels (Methods). The top three lines in the proximity are shown. The circle is colored based on the dendrogram, the mean z-score intensities and the position of the clustered pixels in the PCA cluster image (Methods). The colors of the clustered pixels are assigned on the original IRIS image as a SR cluster image (bottom center). The mean z-score intensity of each target is calculated in each color (bottom right). (**B–E**) Extraction of the red, yellow and green pixels (**B**), the yellow, green and orange pixels (**C**), the red, pale pink and cyan pixels (**D**), and the orange, pale pink, cyan, slate blue, and blue (**E**) in the SR cluster image.

Colocalization was further characterized by the Pearson correlation coefficient between the targets in each colored pixel (Fig. 3). The correlation coefficients between EGFR and Grb2 were 0.79, 0.93 and 0.76 in red, yellow, and green pixels, respectively (Fig. 3A, black arrows). These high correlation coefficients at a 5 nm pixel size, which is close to the size of the protein, strongly indicate the formation of the complex of EGFR and Grb2. In other colors, the correlation coefficient was low. These results indicate complex formation of EGFR and Grb2 in specific regions near the CCS rim (Fig. 2B). We also noted that the correlation coefficient between Grb2 and Eps15 was 0.8 in orange pixels, indicating the complex formation of Grb2 and Eps15 in orange pixels (Fig. 2C and Fig. 3A, H, open arrow), which is not apparent in the original IRIS image (Fig. 1D). The slopes of the regression lines in green, yellow and red pixels were larger in this order (Fig. 3I). This increase in the composition ratio of Grb2 to EGFR toward the center of CCSs suggests gradual complex formation of EGFR with Grb2. PC-coloring in combination with the correlation analysis allows us to identify the sites of complex formation between specific combinations of targets in multiplexed super-resolution imaging.

**Fig. 3.**
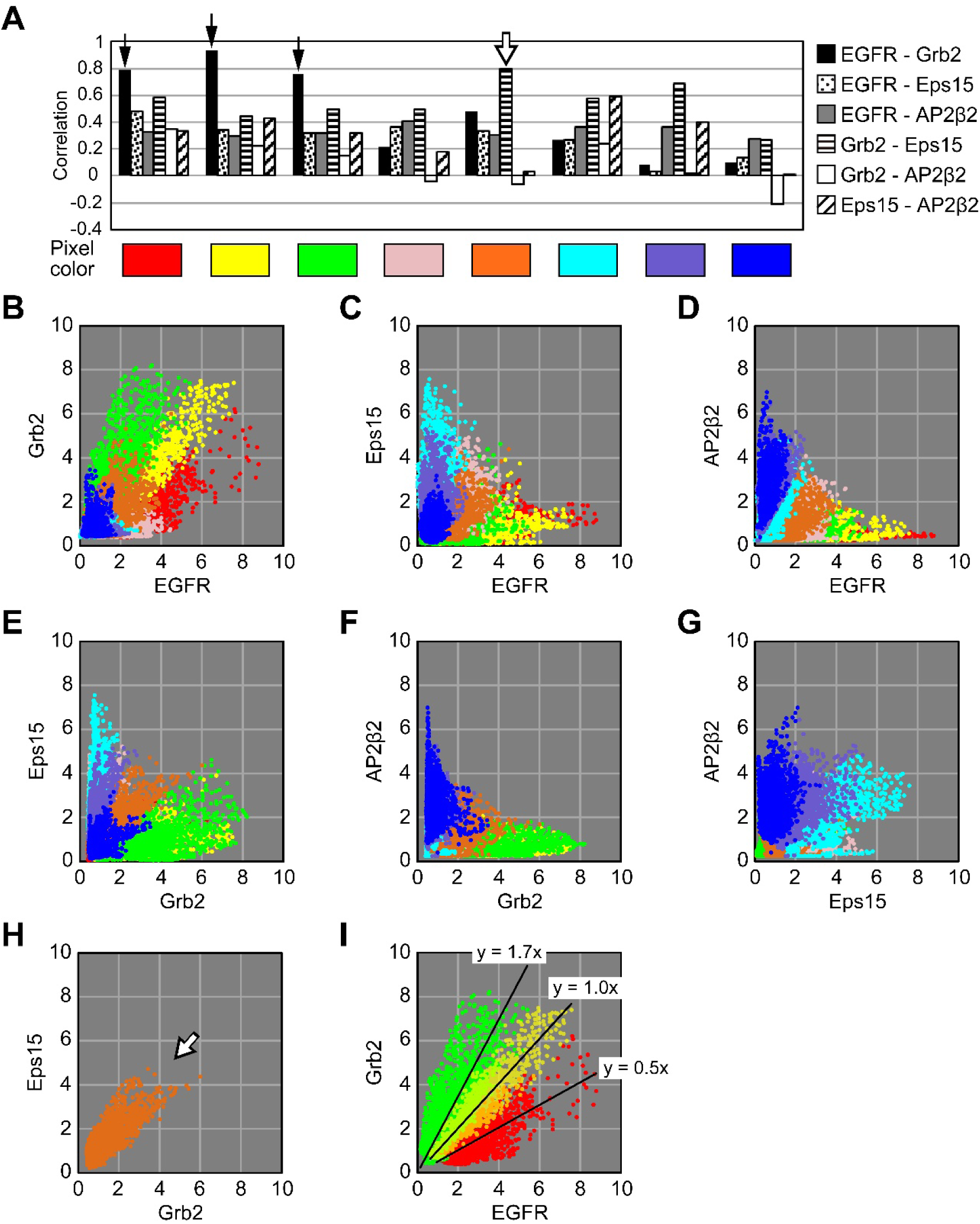
Correlation analysis of the colored pixels shown in Fig. 2. (**A**) Pearson correlation coefficients between their targets. (**B–G**) Scatter plots of z-score intensities of the targets at the pixels in the IRIS images shown in Fig. 2A. The z-score intensities were plotted between EGFR and Grb2 (**B**), Eps15 (**C**) or AP2β2 (**D**), between Grb2 and Eps15 (**E**) or AP2β2 (**F**), and between Eps15 and AP2β2 (**G**). (**H**) Scatter plot of orange pixels between Grb2 and Eps15. (**I**) Scatter plots of red, yellow and green pixels between EGFR and Grb2. Their regression lines are drawn to estimate the composition ratio of Grb2 to EGFR.

## Laminar distribution of the molecular complexes at CCS rims

To further examine colocalization sites of CCS components (Eps15, FCHo1/2 and ITSN1) with EGFR and Grb2, we created SR cluster images (Fig. 4 and Extended Data Figs. 6–8). In the SR cluster image of EGFR, Grb2 and Eps15, EGFR-dominant (orange), EGFR-Grb2-dominant (yellow) and Grb2-dominant (green) pixels lined up from the outside the CCS rim (Fig. 4A). Grb2-Eps15-dominant (magenta) pixels were localized along the edges of Grb2-dominant (green) pixels. On the central side of the CCS, EGFR- Eps15-dominant (cyan) and Eps15-dominant (blue) pixels were localized. Among these color pixels, the yellow, magenta and cyan pixels had a high correlation coefficient of 0.82 between EGFR and Grb2, 0.8 between Grb2 and Eps15, and 0.78 between EGFR and Eps15, respectively, indicating their complex formation (Fig. 4A, red arrows). Eps15 are known to be bound to EGFR upon EGF stimulation^25^. The distribution of cyan pixels indicates that EGFR may form a complex with Eps15 primarily on the central side of the CCS, not in the EGFR clusters at the outer rim of CCSs. Notably, colocalization of Grb2 and Eps15 (magenta pixels) was confined to the boundaries of the Grb2 clusters and the CCS rim. In other SR cluster images, magenta pixels were also confined to the boundaries (Fig. 4B, C). In the scatter plots of Grb2 and Eps15, Eps15 intensity increased with Grb2 intensity at a certain intensity ratio in the magenta pixels (Fig. 4E, arrow). This indicates complex formation of Grb2 and Eps15 with a certain composition ratio in most magenta pixels. On the other hand, in the scatter plots of EGFR and Eps15, the distribution of magenta pixels showed only a partial correlation compared that of the cyan pixels (Fig. 4F, arrows). This suggests that the complex of EGFR and Eps15 is not formed in a large fraction of magenta pixels. This was also seen in the scatter plots of other CCS components (Extended Data Fig. 7E, 8E, right graph, arrows). These results indicate that EGFR recruitment sites at CCS rims are layered from the outside with EGFR-dominant sites, the sites of EGFR-Grb2 complex formation, and Grb2-dominant sites, and the sites of Grb2-CCS component complex formation.

**Fig. 4.**
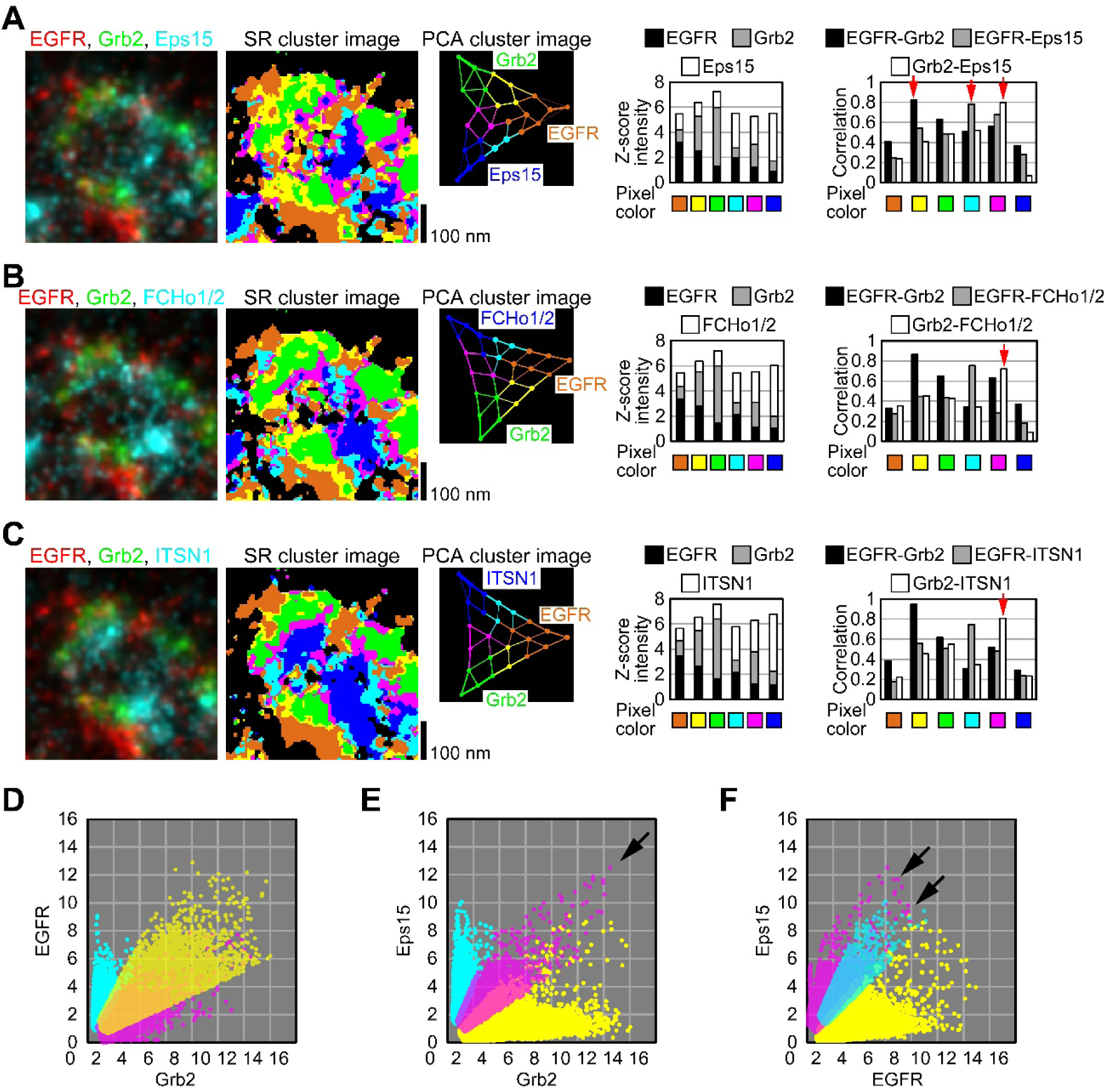
Colocalization analysis of EGFR, Grb2 and Eps15 (A), FCHo1/2 (B) or ITSN1. **(C).** (**A**–**C**) From left to right: the merged IRIS images reconstructed by Gaussian rendering with mean localizations at 5 nm sized pixel, the SR cluster images, the PCA cluster image, the mean z-score intensities of the targets in the clustered pixels and the correlation coefficients between the targets. These PC-coloring with PCA and clustering were performed on the merged IRIS images shown in **Extended Data Figs. 6A, 7A, 8A**. HeLa cells were stimulated with 100 ng/ml EGF for 20 min at room temperature. (**D–F**) Scatter plots of yellow, magenta and cyan pixels shown in **A**. These pixels were plotted between Grb2 and EGFR (**D**), between Grb2 and Eps15 (**E**), and between EGFR and Eps15 (**F**). The scatter plots of other target combination are shown in **Extended Data Fig. 6C, D**.

Next, we examined a spatial relationship among FCHo1/2, Eps15 and ITSN1 (Fig. 5A and Extended Data Fig. 9A). In the SR cluster image, FCHo1/2-dominant (red), FCHo1/2-Eps15-dominant (yellow) and Eps15-dominant (green) pixels were localized at the CCS rim (Fig. 5B and Extended Data Fig. 9B). At the interior of these colored pixels, Eps15-ITSN1-dominant (purple) and ITSN1-dominant (blue) pixels were localized. Among these color pixels, yellow and purple pixels had a high correlation coefficient of 0.73 between FCHo1/2 and Eps15, and 0.78 between Eps15 and ITSN1, respectively (Fig. 5C, arrows). FCHo1/2-ITSN1-dominant (orange) and FCHo1/2-Eps15-ITSN1-dominant (pale pink) pixels also had a high correlation coefficient, but these clustered pixels were small and dispersed (Fig. 5B, C). Overexpression of mCh-FCHo2 highlighted its localization to the CCS rim (Supplementary Fig. 9A). The SR image again shows that the interior of Eps15-FCHo1/2 dominant pixels is lined with Eps15-ITNS1 dominant pixels (Supplementary Fig. 9A). These results indicate that Eps15 colocalizes with different partners from the rim to the center of the CCS in a stepwise manner, which is not apparent in the original IRIS image (Fig. 5A and Extended Data Fig. 9A). In the scatter plots, FCHo1/2 and ITSN1 intensity increased with Eps15 intensity at a certain ratio in yellow and purple pixels, respectively (Fig. 5D, arrows and Extended Data Fig. 10). FCHo1/2, Eps15 and ITSN1 are known to bind to each other^19,26–28^. However, our results suggest that Eps15 forms a complex with FCHo1/2 and ITSN1 in distinct regions. These results reveal the laminar distribution of distinct complexes composed of FCHo1/2, Eps15 and ITSN1 from the rim to the center of CCSs.

**Fig. 5.**
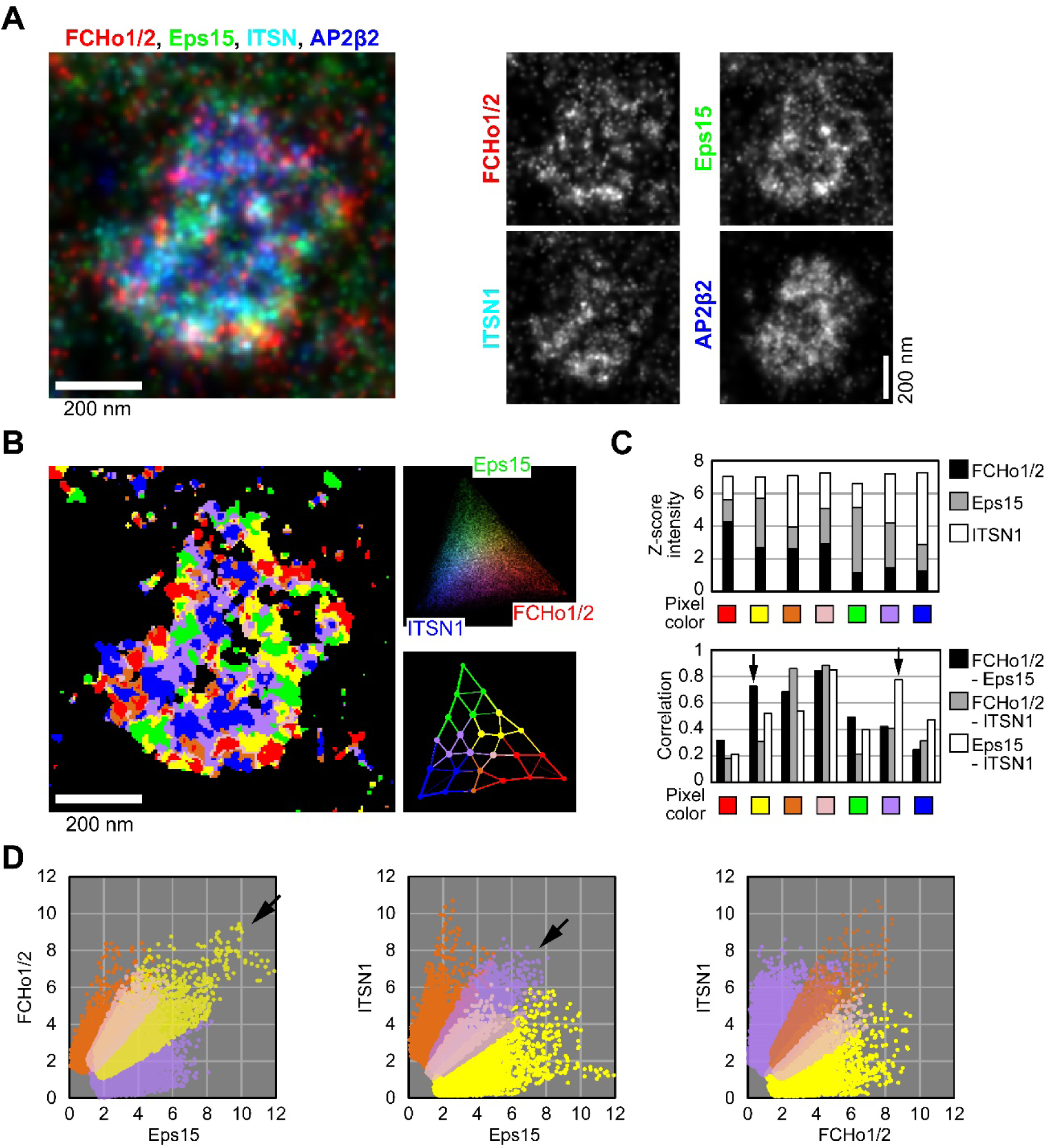
Colocalization analysis of FCHo1/2, Eps15 and ITSN1. (**A**) The merged IRIS image of FCHo1/2 (red), Eps15 (green), ITSN1 (cyan) and AP2β2 (blue) was reconstructed by Gaussian rendering with mean localizations at 5 nm sized pixel. (**B**) The SR cluster image (left), the PCA score image (right upper) and the PCA cluster image (right lower) of FCHo1, Eps15 and ITSN1. (**C**) The mean z-score intensities of the targets in the clustered pixels (upper) and the correlation coefficients between the targets (lower). These PC-coloring with PCA and clustering were performed on the merged IRIS images shown in **Extended Data Fig. 9A**. (**D**) Scatter plots for comparing FCHo1/2-Eps15 dominant (yellow), FCHo1/2-ITSN1 dominant (orange), FCHo1/2-Eps15-ITSN1 dominant (pale pink) and Eps15-ITSN1 dominant (purple) pixels. The scatter plots of other target combination are shown in **Extended Data Fig. 10**.

## Recruitment sites of membrane receptors

We compared the recruitment sites of EGFR and TfR in a CCS. In the merged IRIS image, TfR clusters were localized in spots (Fig. 6A, cyan arrowheads). The SR cluster image showed that TfR-dominant pixels (cyan) were separated from EGFR-dominant (red) and Grb2-dominant (green) pixels (Fig. 6B). In the scatter plots of TfR and EGFR or Grb2, the TfR-dominant pixels (cyan) were largely separated from both EGFR-dominant (red) and Grb2-dominant (green) pixels (Fig. 6E, center and right, arrows), compared to the position of the red and green pixels in the scatter plot of EGFR and Grb2 (Fig. 6E, left, arrows). Since EGFR and TfR are expected to accumulate at the endocytic sites, the separation of these dominant pixels suggest that their endocytic sites are different.

**Fig. 6.**
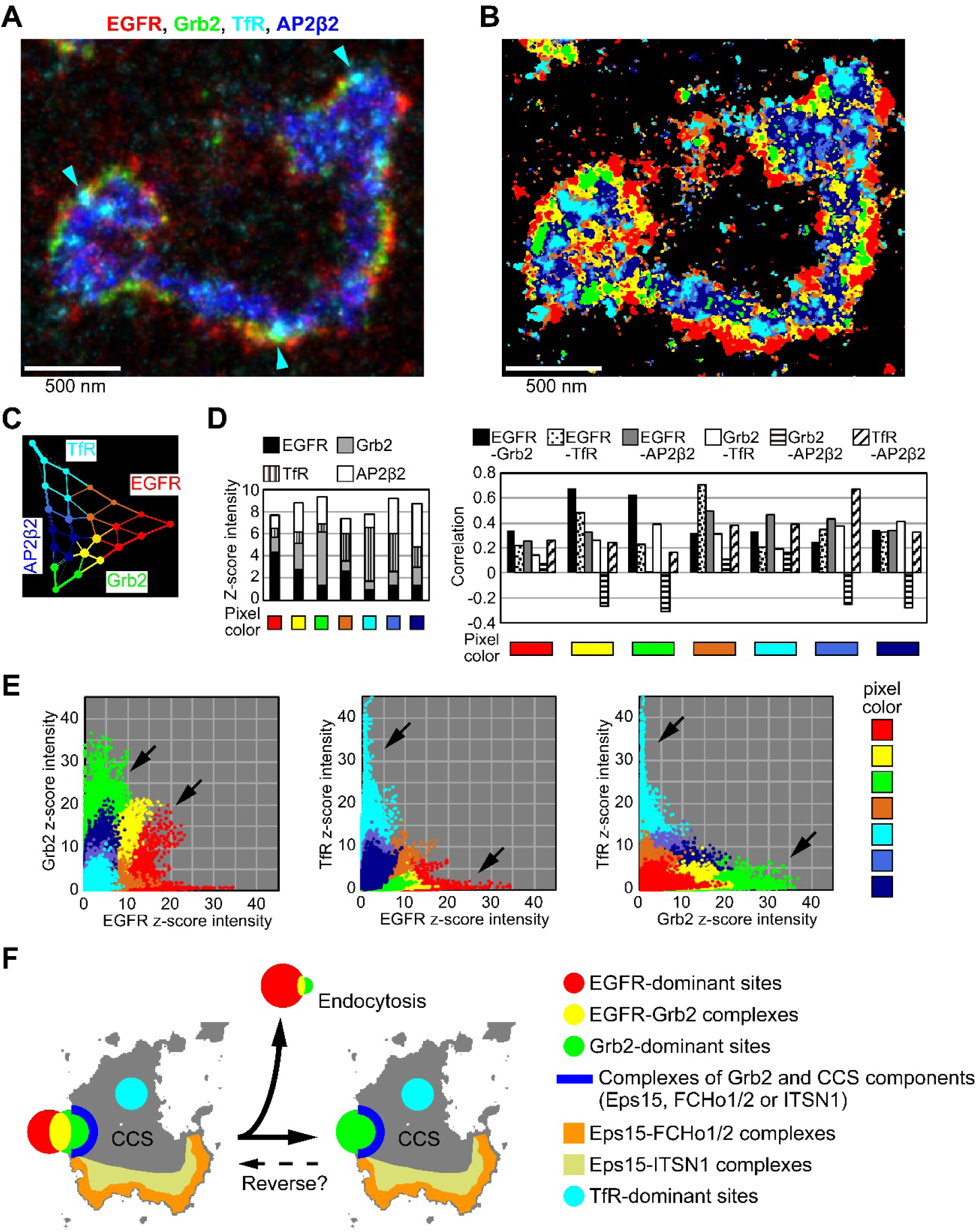
Colocalization analysis of EGFR, Grb2, TfR and AP2β2. (**A**) Merged IRIS image of EGFR (red), Grb2 (green), TfR (cyan) and AP2β2 (dark blue) was reconstructed by Gaussian rendering with mean localizations at 5 nm sized pixel. (**B**, **C**) The SR cluster image (**B**) and the PCA cluster image (**C**) of EGFR, Grb2, TfR and AP2β2. (**D**) The mean z-score intensities of the targets in the colored pixels (left) and the correlation coefficients between the targets (right). The IRIS images including 112 CCSs were subjected to PCA and cluster analysis with the same eigenvectors and clustering. The cells were stimulated with 100 ng/ml EGF for 20 min at room temperature or for 3 min at 37℃ in total three independent experiments. (**E**) Scatter plots of z-score intensities of the targets at the pixels in the IRIS images of the 112 CCSs. The z-score intensities were plotted between EGFR and Grb2 (left) or TfR (center), and between Grb2 and TfR (right). (**F**) The laminar complex formation of EGFR, Grb2 and CCS components at EGFR recruitment sites in CCS rims.

## Discussion

Colocalization analyses using single-molecule localization-based super-resolution microscopy have been reported for Clus-Doc^29^, ClusterViSu^30^ and Coloc-Tesseler^31^, but these analyses are performed on only two targets. In this study, we developed PC-coloring for colocalization analysis of three or more targets. In the first step, PC-coloring creates the color map of colocalization sites characterized by the intensity ratio of the targets. In the second step, the detailed correlation between individual target pairs is examined in the scatter plots. A high correlation coefficient may reflect the formation of the molecular complexes because correlation is analyzed with highly accurate quantitative data in protein-size pixels. Hence, in PC-coloring, PCA extracts features about molecular complexes from the multiplexed intensity data. By assigning the features on the original image, their spatial relationship is visualized. PC-coloring was enabled by the advantages of IRIS. The high-density labeling with the antiserum-derived Fab probes allowed faithful visualization of distributions of multiple endogenous proteins. This high density also allowed improved resolution using the large number of probes bound over multiple frames. These resulted in greater separation of the pixels by PCA.

At the CCS rim, the localizations of various endocytic proteins have been reported^16,19,20^. In addition, many clathrin-coated microcages of 50 – 60 nm diameter emerge from CCS rims of cells treated with a proton ionophore at pH 6.3 ^32^, similar to pH inside endocytic vesicles containing EGF^33^. Therefore, the CCS rim has been considered an endocytic site of receptor^13,20^. EGF-induced colocalization of EGFR, Grb2 and CCS components shown in this study provides direct evidence for molecular complexes at the EGFR recruitment sites in the CCS rim. The separation of EGFR- dominant and Grb2-dominant regions from TfR-dominant regions suggests EGFR-specific endocytosis sites at the CCS rim. This is consistent with the facts that Grb2 is required for endocytosis of EGFR, but not TfR^34^, and that the sorting of EGFR and TfR occurs early in the endocytosis process on the CCSs^35,36^. Furthermore, the laminar complexes of EGFR, Grb2 and CCS components suggests that Grb2 anchors EGFR to CCS rims. This is supported from the domain structure of Grb2, in which its SH2 domain binds to phosphorylated EGFR, and its two SH3 domains bind to the proline-rich domains of Eps15 and Dynamin^37,38^. Indeed, overexpression of the SH3 domain-deletion mutant of Grb2 inhibits EGFR recruitment to CCSs^14^. The half-life of Grb2 binding to EGFR is about five sec^39^, which is shorter than the endocytosis process of 1 – 2 min^40^. Thus, at the EGFR recruitment sites, Grb2 can form a complex with CCS components that remains after EGFR endocytosis.

FCHo1/2, Eps15 and ITSN1 were reported to be localized around the rim of the CCSs^16,19,20^. This study further clarified that the interior of Eps15-FCHo1/2 complexes are lined with Eps15-ITSN1 complexes at the CCS rims. These proteins can form homo- oligomers^41–43^. The F-BAR protein, FCHo1/2, senses membrane curvature and binds to the plasma membrane^42^. Eps15 is strongly bound to FCHo1/2 because it is deprived of CCSs by the ectopically localized C-terminal domain of FCHo1/2^19^. Furthermore, Syp1p and Ede1p, yeast homologs of FCHo1/2 and Eps15, are the earliest to assemble at the endocytosis site in yeast, followed by Pan1p and Sla1p, homologs of ITSN1^44^. Together with these reports and our results, Eps15 clusters may grow from Eps15-FCHo1/2 complexes, and then ITSN1 clusters may grow on the Eps15 clusters as a foothold in CCS formation.

Our results indicate six layers at EGFR recruitment sites in CCS rims (Fig. 6F), as follows: (i) EGFR-dominant sites, (ii) the sites of EGFR-Grb2 complex formation, (iii) Grb2-dominant sites, (iv) the sites of Grb2-CCS component complex formation, (v) the sites of Eps15-FCHo1/2 complex formation, (vi) the sites of Eps15-ITSN1 complex formation. After EGF endocytosis, the remaining Grb2 clusters could trap another EGFR cluster. The colocalization of SOS and cCbl suggests that this laminar complex is also the initiation site of signal transduction. The signaling would be enhanced in many tumor cells in which endocytosis is delayed by overexpression or heterodimers of EGFR^45–48^. PC-coloring can update the *in vitro* network of protein interactions^15,49,50^ to the *in vivo* map of protein complexes.

## METHODS

### Plasmids and reagents

Expression plasmids for human CLC (#57452), human TfR (#61505), human Eps15 (#27696), human ITSN1 Long (#47395), mouse FCHo2 (#27686), human Epsin2 (#22242), mouse Hip1r (#), human Dynamin2 (#27689) and human AP2μ2 (#27672) were purchased from Addgene. cDNA encoding human AP2β2 (Clone ID: 3532472) and human FCHo1 (Clone ID: 5218319) were purchased from Horizon (Dharmacon). cDNA encoding nanobodies for anti-EGFP and anti-mCh were synthesized by gBlocks Gene Fragments (Integrated DNA Technolodies) according to reported sequences (Clone ID: LaG-3 and LaM-1)^24^. Expression plasmids for human EGFR, human cCbl and human Grb2 were previously described^48,51^. These cDNAs and their fragments were inserted into pGEX-6P-1 vector using PCR for protein purification from E. coli. EGF was purchased from Pepro Tech. DyLight 488 NHS Ester was purchased from Thermo Fisher Scientific.

### Cell culture

HeLa cells were cultured in DMEM supplemented with 10% fetal bovine serum. For stimulation with 100 ng/ml EGF, the cells were serum-starved overnight. For plasmid transfection, FuGENE HD (Promega) was used, and the cells were cultured for 24 hours before IRIS imaging.

### Purification of antigen proteins and nanobodies

In this study, 4 to 5 mg of a recombinant protein per a Fab probe was required for rabbit antiserum preparation, the affinity purification of polyclonal antibodies and Fab probe purification. To increase the yield of the recombinant protein, a general purification protocol for glutathione S-transferase (GST)-fused protein from E. coli was improved using an excess of glutathione resins. BL21(DE3) competent cells were transformed with pGEX-6P-1 vector encoding GST-fused protein. After 35 μM IPTG induction, 6L cultures of the cells were incubated for 16 to 20 hours at 20℃ with shaking. The collected cells were lysed in 50 ml lysis buffer (20 mM Tris pH7.3, 150 mM NaCl, 0.25% Triton X-100, 1 mM DTT) containing protease inhibitor cocktail (Nacalai Tesque) and sonicated. Glutathione resins (GST-Accept, Nacalai Tesque) in 30 to 60 ml bed volume was added to the supernatant of the cell lysate, and then was rotated for 90 min at 4 ℃. The resins were transferred to two or three columns constructed from a 50-ml plastic syringe. The resins were washed with the lysis buffer and a high Na buffer (20 mM Tris pH7.3, 500 mM NaCl, 0.25% Triton X-100, 1 mM DTT), and three times for 5 min with a buffer A (20 mM Tris pH7.3, 500 mM NaCl, 5 mM MgCl, 0.25% Triton X-100, 1 mM DTT, 1mM ATP). The resins were further washed five times with a PreScission reaction buffer (50 mM Tris pH7.3, 150 mM NaCl, 1 mM EDTA, 1 mM DTT) for antiserum preparation. For affinity column preparation, the resins were washed five times with a NaHCO_3_ buffer (200 mM NaHCO_3_, 500 mM NaCl) to remove primary amines from the buffer. Two ml of 50% slurry of the resins were aliquoted into thirty to sixty 2-ml tubes. The resins were subject to PreScission protease and rotated overnight at 4℃ to elute the target proteins. The use of the 2-ml tubes is critical in this elution step. For unknown reasons, some proteins such as FCHo1, Dyn2 and cCbl were reproducibly lost in this step when 50-ml or 15-ml tubes were used, even if low-protein adsorption tubes (PROTEOSAVE, Sumitomo Bakelite Co.) were used. The eluted protein solution was in turn transferred to an ultrafiltration column (Amicon Ultra-15, Merck Millipore) and concentrated by centrifugation. Eighty to ninety percentage of the eluted protein was recovered from the column, if the water level of the solution was not lowered below the height of the column’s filter during centrifugation. The resins were again filled with the buffer to obtain the second and third eluates, which were added to the first eluate and all concentrated to 2 ml. Since the predicted molecular weights of TfR1-60, TfR93-179, Grb2 109-159 and Grb2 160-217 are small (6–10 kDa), for antiserum preparation, the GST-fused protein fragments were eluted with 20 mM reduced glutathione. The reduced glutathione was removed by dialysis. By this protocol using an excess of glutathione resins, the yield of EGFR 985-1210, which had a low yield, increased about 11-fold. Seventeen antigens, some with molecular weights around 100 kD, were obtained in milligram quantities in one or two experiments, for FCHo2 in three or four experiments (Supplementary Table 2). Nanobodies for EGFP and mCh were purified using the usual amount of the glutathione resins from BL21 cells because of their high yield.

The used resins were collected and reused after being cleaned according to the cleaning protocol for a glutathione column (GSTrap^TM^ FF, Cytiva). The resins were transferred to two or three columns constructed from a 50-ml plastic syringe. GST remained on the resins was removed by adding 20 mM reduced glutathione three times. The resins were washed five times with PBS. Precipitated substances were removed with 6 M guanidine hydrochloride (denaturing agent). The resins were washed fifteen times with PBS. Hydrophobically bound substances were removed by adding 70% ethanol three times. The resins were stored in 20% ethanol at 4℃. No protein bands were detected when the cleaned resins were subjected to SDS–polyacrylamide gel electrophoresis.

### Production of antiserum-derived Fab probes

IgG antibodies in a rabbit antiserum were purified by a Protein A column (HiTrap rProteinA FF 5 ml, Cytiva). After dialysis with PBS, the IgG antibodies was applied to an affinity column (HiTrap NHS-activated HP 1 ml, Cytiva) coupled to 1.5–3 mg of its antigen protein. For IgG antibodies from antiserum against GST-tagged TfR 1-60, TfR 93-179, Grb2 109-159 or Grb2 160-217, the affinity column was coupled to GST-cleaved TfR 1-60, TfR 93-179 or Grb2 full length, respectively. In addition, the column was linked to a GST-coupled affinity column to remove anti-GST antibodies. The polyclonal antibodies were eluted with 100 mM glycine buffer at pH 3.5 (or 3.0) and pH 2.0. The pH of the eluate was immediately neutralized with 1 M Tris buffer at pH 9.0, and then dialyzed in PBS. To find the eluted fraction that can stain for endogenous targets, HeLa cells were immunostained. For fluorescent labeling, the fraction was added to a 200 μl bed volume of antigen-conjugated resins (NHS-activated Sepharose 4 Fast Flow, Cytiva) to protect their antigen binding sites. The antigen bound to the resins was more than twice the molar amount of the antibodies in the fraction. After overnight rotation at 4℃, the antibodies were labeled with twice the molar amount of Dylight 488 NHS Ester as the total amount of the antigens and antibodies on the resins. After rotation at room temperature for 1 hour, the resins were transferred to a low-protein adsorption 1.5-ml tubes (PROTEOSAVE, Sumitomo Bakelite Co.), whose bottom was perforated with a fine needle. The tube was placed on top of a 2-ml tube to construct a homemade microspin column. The resins were washed five times with PBS. The antibodies were eluted three times with 200 μl of 100 mM glycine buffer at pH 2.0. The pH of the eluate was immediately neutralized with 1 M Tris buffer at pH 9.0. The eluted antibodies were mixed with the first and second eluates, and then digested with papain to generate Fab fragments. Their Fc domains were removed using Protein A Sepharose CL-4B (Cytiva). To reduce the non-specific signals of the Fab, the Fab was further bound to antigen-conjugated resin. After overnight rotation at 4℃, the resins were transferred to the handmade microspin column constructed from a 0.5-ml tube and a 1.5-ml tube. The resins were washed five times each with 100 μl of PBS and 100 mM glycine buffer at pH 5.0. The Fabs were eluted ten and five times with 100 μl of 100 mM glycine buffer at pH 3.0 and pH 2.0, respectively. The pH of the eluate was immediately neutralized with 1 M Tris buffer at pH 9.0. The labeling ratio of the Fabs was estimated to be about 0.6 by enzyme-linked immunosorbent assay (ELISA). The fraction eluted at pH 3.0, which contains the highest amount of the Fabs, was used for IRIS super-resolution imaging. In each of these purification steps, the polyclonal antibodies and Fabs were placed in the low-protein adsorption tubes to protect them from being lost. This protocol was established using anti-CLC antiserum (Supplementary Fig. 1).

### Multiplexed super-resolution imaging

Multiplexed IRIS super-resolution imaging was performed as previously described^6^. HeLa cells on a 24 mm round coverslip were fixed with 3.7% PFA in a cytoskeleton buffer (10 mM MES pH 6.1, 90 mM KCl, 3 mM MgCl_2_, 2 mM EGTA) for 20 min, and then permeabilized with 0.2% Triton-X100 in a HEPES buffered solution (10 mM pH 7.3, 90 mM KCL, 3 mM MgCl_2_, 100 μM DTT) for 5 min. After blocking with 4% bovine serum albumin in the HEPES buffered solution for 30 min, the coverslip was mounted in an open chamber (A7816, Thermo Fisher Scientific). On the coverslip, a handmade chamber with a volume of approximately 50 μl was constructed with a vacuum grease and a 1 cm diameter plastic ring. As a retractable lid, a 15 mm round coverslip was placed on top of the ring and another 24 mm round coverslip was placed on top of the open chamber to double the lid. The cells were subjected to the IRIS probe in an imaging solution (50 mM Tris pH 8.0, 90 mM KCl, 3 mM MgCl_2_) with an oxygen scavenging mix (5 U/ml pyranose oxidase, 10 mM cysteamine hydrochloride, 60 μg/ml catalase, 10% glucose)^5^. The probe concentrations in the imaging solution, measured as the concentration of the conjugated fluorescent dye (DyLight 488), were 80 nM anti-EGFR Fab, 40 nM anti-CLC Fab, 100 nM anti-Eps15 Fab, 40 nM anti-TfR Fab, 80 nM anti- Grb2 Fab, 30 nM anti-AP2β2 Fab, 60 nM anti-ITSN1 Fab and 60 nM anti-FCHo1 Fab. The single molecule speckle (SiMS) images were acquired by TIRF microscopy using an inverted microscope (Olympus IX83-ZDC) equipped with an IX3-ZDC2 Z-drift compensator (Olympus), an UPlanApo 60×/1.50 NA oil objective lens (Olympus), ORCA-Flash4.0 V3 Digital CMOS camera (Hamamatus) and a 488 nm laser line (150 mW, Coherent). A set of a bright field image (exposure time, 500 ms) and 500 SiMS images (exposure time, 100ms) was acquired repeatedly. The bright field images were used to correct for stage drift (see below). The total number of SiMS images acquired was as follows: 250,000 (anti-EGFR Fab), 100,000 (anti-CLC Fab), 300,000 (anti-Eps15 Fab), 200,000 (anti-TfR Fab), 400,000 (anti-Grb2 Fab), 200,000 (anti-AP2β2 Fab), 400,000 (anti-ITSN1 Fab) and 400,000 (anti-FCHo1 Fab). After IRIS imaging of each probe, the probe was washed out 15 times with 0.2% Triton-X100 in the HEPES buffered solution and 10 times with the HEPES buffered solution. The remaining fluorescence of the probe was completely photobleached in the HEPES-buffered solution with the oxygen scavenging mix.

### Image reconstruction

In hundreds of thousands of the SiMS images, the 10^6^ to 10^8^ localizations of fluorescent speckles were determined using ThunderSTORM plugin^52^ in ImageJ (http://rsb.info.nih.gov/ij/). Stage drift during long-time image acquisition for each probe was corrected by an autocorrelation function using the bright field image of fixed cells and non-fluorescent beads attached to the coverslip as previously described^6^. After the drift correction for each probe, stage drift between two different probes was corrected by calculating cross-correlation functions between 30 bright field images per probe. The 30 × 30 cross-correlation functions were computed using the 10-fold magnified images with a bicubic method (pixel size, 10.83 nm). The drift distances between the two probes were calculated, and its standard error was around 1 nm.

To improve the localization accuracy, the mean localization and standard error of a probe bound over two frames were used. The localizations of a long-binding probe were distributed within a radius of 54 nm. The duration of the quenching state due to blinking was one frame (exposure time: 100 ms). Thus, localizations that were continuous or one frame apart within a radius of 54 nm were extracted as the localizations of the same probe bound over two frames. To reduce the contribution of the first localization as a reference, the mean of the extracted localizations was calculated. Using the mean localization as a new reference, the localizations were re-extracted using the same filter. The mean and standard error of the re-extracted localizations were calculated. These calculations were computed using our plug-in with multi-threaded processing in ImageJ. In addition, the accuracy of the drift correction using the bright field image was estimated from differences between the drift-corrected localizations of the same probe in two SiMS images acquired before and after the bright field image. The mean of the difference was about 3 nm. Together with the accuracy of the drift correction between the two different probes, the standard error of the localizations less than 5 nm was converted to 5 nm. Using the mean localization and the standard error, an IRIS image was reconstructed at 5 nm pixel size by using a Normalized Gaussian rendering in the ThunderSTORM. When Gaussian rendering was not used, the IRIS image was reconstructed by integrating the localizations as previously described^6^.

### PC-coloring

PC-coloring was computed using a set of our plug-ins in ImageJ and scripts in Matlab (MathWorks). Each process using the ImageJ plug-ins or Matlab scripts is indicated at the end of the following sentences as (ImageJ) or (Matlab), respectively.

A data set of the intensities of each target and the coordinates of its pixel in the IRIS images was output (ImageJ). In the multiplexed intensity data, a pixel with target intensities of all 0’s was excluded. To match the intensity distribution among targets, the intensity data was standardized (z-score) by each target (Matlab). To calculate the intensity ratio at each pixel, if this z-score intensity data had a value less than 0, an appropriate value was added to all the intensities to make them greater than or equal to 0 (Matlab). Using the intensity ratios, PCA was performed, and PCA scores were output (Matlab). A binarized image was created from the summed IRIS image of each target to exclude pixels with low intensity (ImageJ). Using the binarized image, the pixels with low intensity were also removed from the PCA score data (ImageJ). To interpret the PCA result, a PCA score image was created by plotting the raw intensity of each IRIS image on the coordinates of the first and second PCs (ImageJ). Then the PCA score images of each target were merged (ImageJ). The PCA score data were clustered by k-means method (Matlab). The number of clusters set by the k-means method was 25 in this study. The similarity of the intensity ratio between the clustered pixels was calculated as the distance by group average method (ImageJ). The two clustered pixels with the closest similarity were combined, and then the group average distance between the new clustered pixels and the other clustered pixels was calculated for their linkage distances (ImageJ). Using the linkage distances, a dendrogram was created (Matlab). The number of the clustered pixels was renumbered in order from left to right of the leaf nodes in the dendrogram (Matlab). The mean z-score intensities of each target in the cluster pixels were output (ImageJ).

A SR cluster image was created by assigning the number of the clustered pixels to the pixel in the original IRIS image (ImageJ). To examine the adjacent clustered pixels on the SR cluster image, 8-connectivity analysis was performed^53^. One pixel in the clustered pixels in the SR cluster image was selected, and the cluster numbers of the surrounding eight pixels were obtained (ImageJ). If any of the cluster numbers of the eight pixels is different from that of the center pixel, the cluster number was recorded as the pixel located at the border of the adjacent cluster (ImageJ). By performing this process on all pixels in the clustered pixels, the cluster numbers of the boundary pixels were tallied, and their percentages were output as proximity between the clustered pixels (ImageJ). A boundary pixel without any cluster number were excluded from this percentage.

A PCA cluster image was created by drawing circles on the coordinates of the first and second PCs (ImageJ). The center of the circle was that of the clustered pixels. The area of the circle was proportional to the number of pixels in the clustered pixels. The two circles were connected by a line with a width proportional to the proximity between the clustered pixels on the SR cluster image, generating a PCA cluster image with proximity (ImageJ). In the PCA cluster image with proximity, the top three lines in their proximity were shown. The sum of the three proximity, on average, exceeds 70% of all proximity. In the SR cluster image, adjacent clustered pixels were also adjacent in the PCA cluster image. This indicates that the adjacent relationship of the clustered pixels in the SR cluster image is also preserved in the PCA cluster image.

The clustered pixels in the SR cluster image and the PCA cluster image with proximity were colored (ImageJ). The clustered pixels were colored based on the dendrogram, the mean z-score intensity of target, and the position of the clustered pixels in the PCA cluster image (ImageJ). The clustered pixels with high similarity in intensity ratio at adjacent positions in the PCA cluster image were colored the same. The coloring was adjusted to reflect the distribution of the target intensities on the PCA score image (ImageJ). In each color pixels, the z-score intensity of the target was plotted on a scatter plot. The Pearson correlation coefficient was calculated (ImageJ). In PC-coloring, there is no principle limit to the number of targets. However, pixel separation by PCA decreases with the number of targets. In this study, three to four targets were practical.

## ACKNOWLEDGMENTS

This work was supported by JSPS KAKENHI Grant Number 24H01282, 21K06168, 19H04961, 17K07384, 15H01635 (T.K.) and 22H00456 (N.W.), by the Asahi Glass Foundation, Takeda Science Foundation (T.K.), Uehara Memorial Foundation (N.W.) and by CREST grant number JPMJCR15G5 (N.W.).

## AUTHOR CONTRIBUTIONS

T.K. improved a method for protein purification from E. coli and developed a method for Fab probe production from antiserum, image reconstruction and PC- coloring. T.K. designed and conducted experiments including protein purification, Fab probe production and microscopic imaging. T.K developed all image analysis programs and analyzed the data. T.K. supervised the project and wrote the manuscript with support from N.W. R.K. contributed to the production of Fab probes for CLC and Eps15, and nanobodies for EGFP and mCherry, constructed expression plasmids and purified proteins. S.O. and L.E. purified proteins.

---

**Extended Data Fig. 1.**
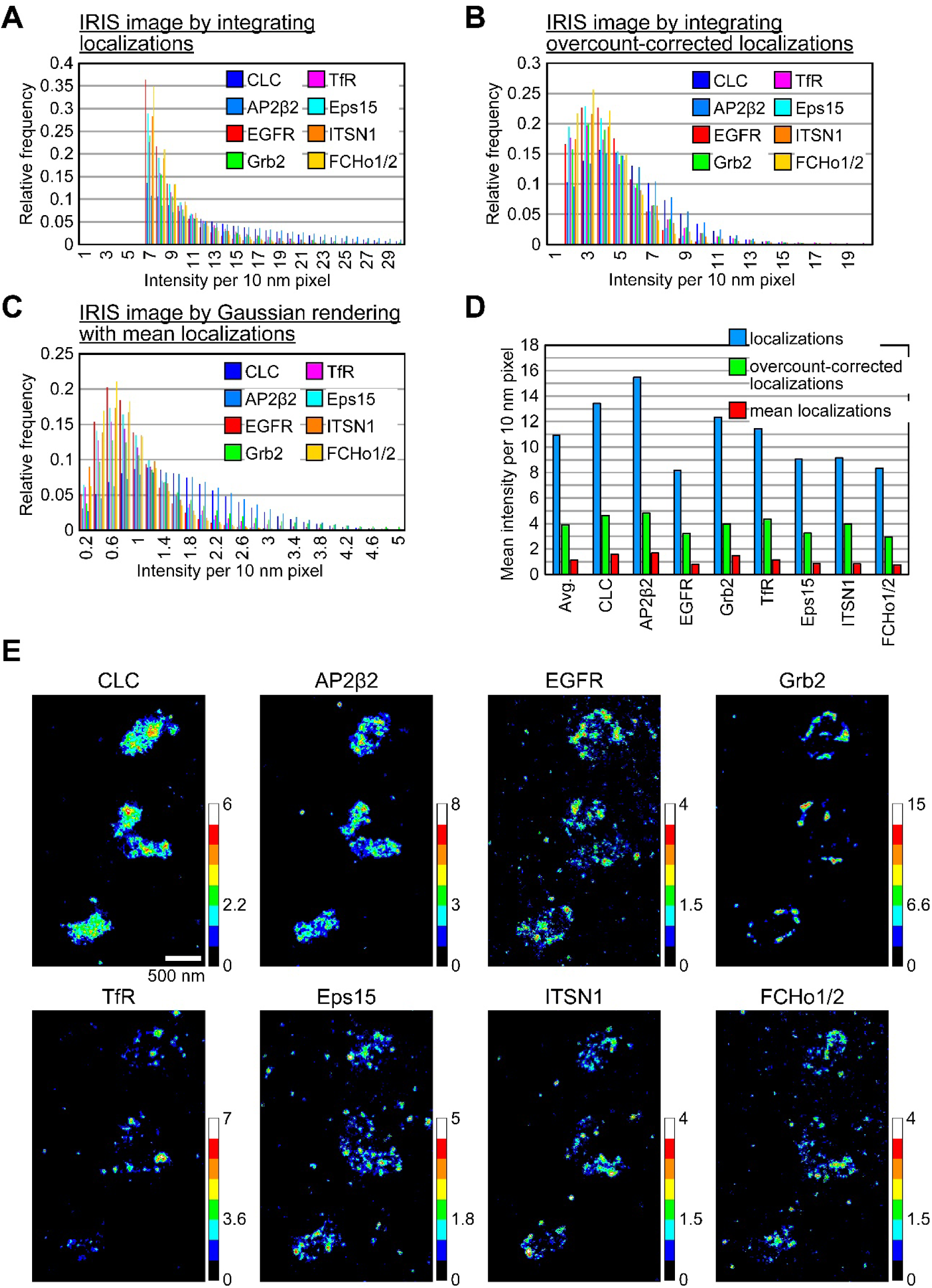
Labeling densities of Fab probes. (**A–C**) Histogram of the raw intensities per 10 nm pixel in the IRIS image reconstructed by integrating localizations (A) and overcount-corrected localizations (**B**) and by Gaussian rendering with mean localizations (**C**). The number of localizations is overcounted more than the number of probe binding events due to probes bound over multiple frames. Therefore, the probes bound over multiple frames were extracted, and their mean localization was calculated as one localization (Methods). The overcounted localizations was corrected by combining the mean localization data with the localization data of once-bound probe. Therefore, in IRIS image reconstructed by integrating the overcount-corrected localizations, the intensity per pixel represents the true density of the bound probes. To measure the intensities per 10 nm pixel in their target-localized regions, a binarized image was created by thresholding an intensity in the IRIS image reconstructed by integrating localizations at 10 nm pixel size. Using the binarized image, the regions with low target intensity were masked in the IRIS image. The intensity per 10 nm pixel in the masked IRIS image was plotted on a histogram. (**D**) Mean intensity per 10 nm pixel of each target in the masked IRIS image. The mean density of the overcount-corrected localizations (i.e. the true density of bound probes) was 3.9 labels per 10 nm pixel, which is 4.4 times the maximum density at which 12 nm sized antibodies can physically bind. In the Gaussian rendering IRIS image, only probes bound over two frames were used, so the intensity per 10 nm pixel was about one-quarter of that in the overcount-corrected IRIS image. (**E**) The masked IRIS images reconstructed by Gaussian rendering with mean localizations in 8 pseudo color. In the target-localized regions, the intensity per 10 nm pixel was 1.5 – 6.6 or higher. These localization data is from the probes in Fig. 1A.

**Extended Data Fig. 2.**
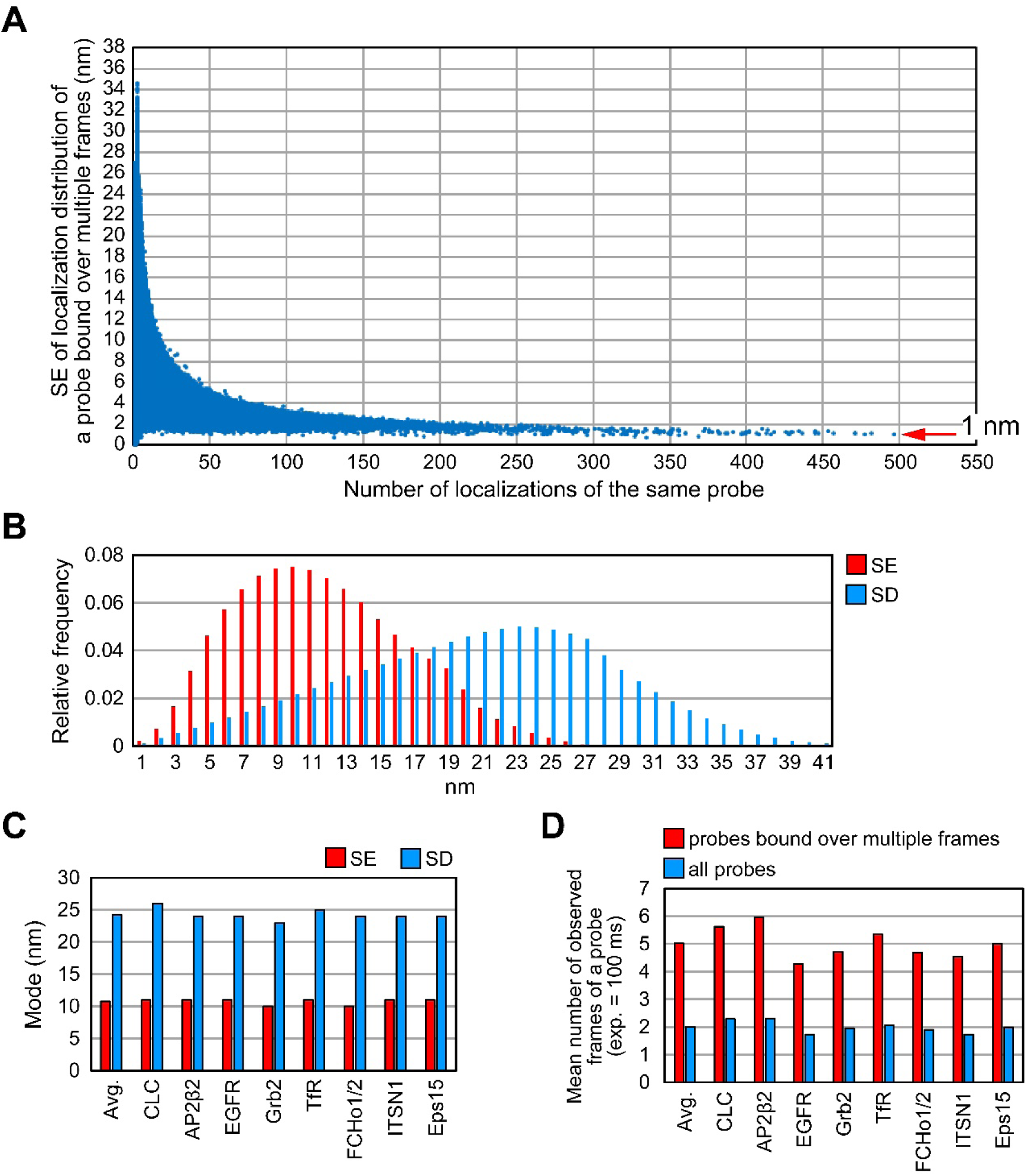
Localization accuracy of Fab probes. (**A**) Scatter plot of standard errors (SEs) of localizations of a probe bound over multiple frames. The localizations of a probe bound over two or more frames were extracted, and the mean localizations and SE were calculated for each probe (Methods). The SE indicates the accuracy of mean localizations and estimates the resolution of the Gaussian rendering image. The SE decreased with the number of the localizations and reached around 1 nm. (B) Histogram of the SE and standard deviations (SD) of the localizations. The SD was calculated from the localization distribution of a probe bound over multiple frames. The SD indicates the spread of the localizations of the same probe and estimates the resolution of the IRIS image by integrating the localizations. The localization data in **A**, **B** is from anti-Grb2 probe in Fig. 1A. (**C**) The mode of the SEs and SDs from the eight probes. On average, the mode of the SE, 10.7 nm, was smaller than that of the SD, 24.2 nm, indicating 2.26-fold improvement in the resolution. (**D**) Mean number of observed frames of a probe. The localization data in **C**, **E** is from the probes in Fig. 1A.

**Extended Data Fig. 3.**
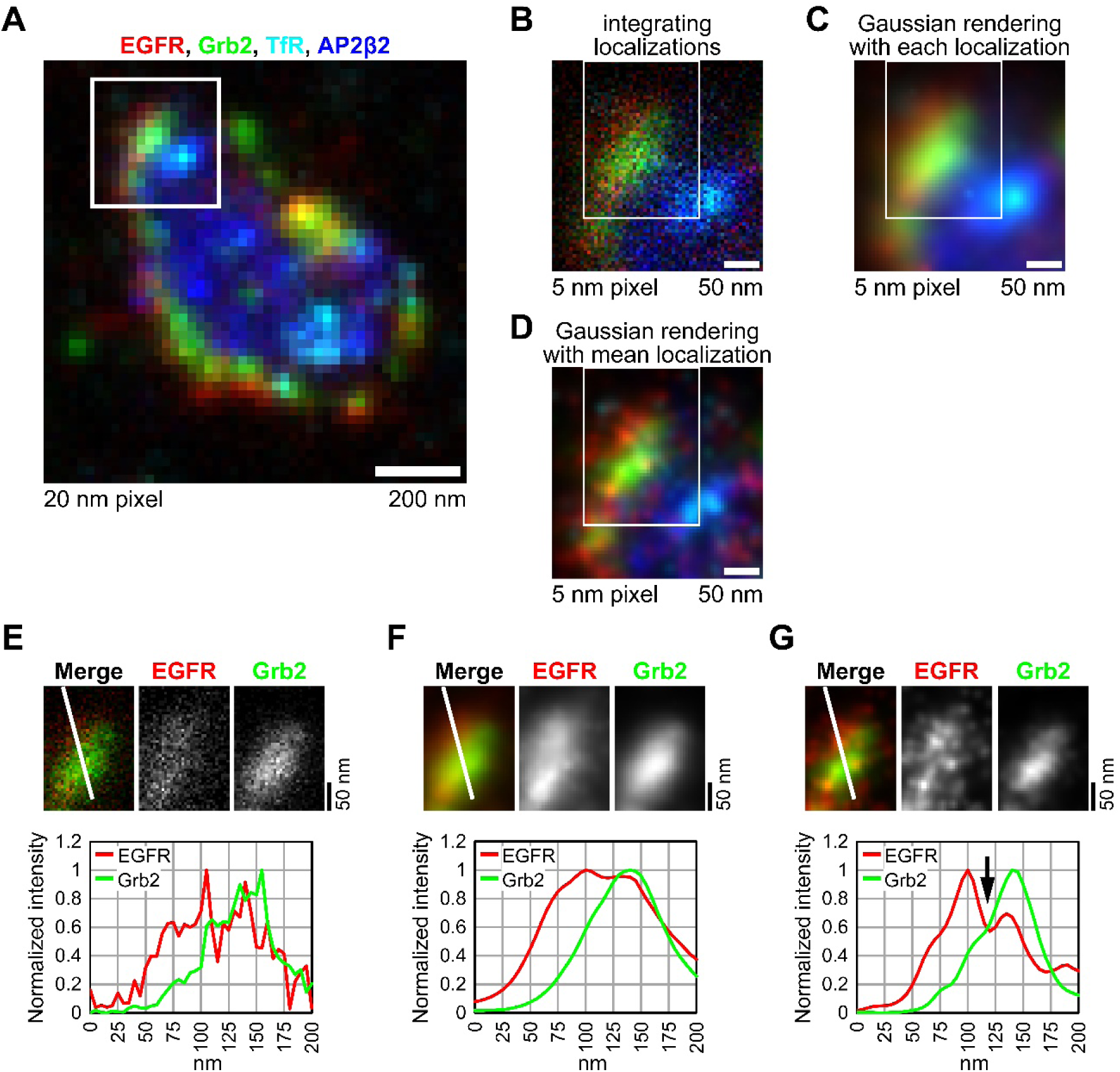
Improvement of the resolution. (**A**) Merged IRIS image of EGFR (red), Grb2 (green), TfR (cyan) and AP2β2 (blue) in the boxed area shown in Fig. 1A. The IRIS image was reconstructed by integrating localizations at 20 nm sized pixel. (**B**) The IRIS image was reconstructed by integrating localizations at 5 nm sized pixel in the boxed area in **A**. (**C**) The IRIS image was reconstructed by Gaussian rendering with each localization and SE at 5 nm sized pixel in the boxed area in **A**. Each localization and SE were calculated by Gaussian fitting of a fluorescent speckle. (**D**) The IRIS image was reconstructed by Gaussian rendering with mean and SE of localizations at 5 nm sized pixel in the boxed area in **A**. The localizations of a probe bound over two or more frames were extracted, and their mean localizations and SE were calculated (Methods). (**E–G**) Cross-sectional profiles of EGFR and Grb2 in the boxed area in **B** (**E**), in **C** (**F**) and in **D** (**G**). The IRIS image by Gaussian rendering using mean localizations reveals that the location of EGFR and Grb2 clusters were shifted at the CCS rim (**G**, arrow).

**Extended Data Fig. 4.**
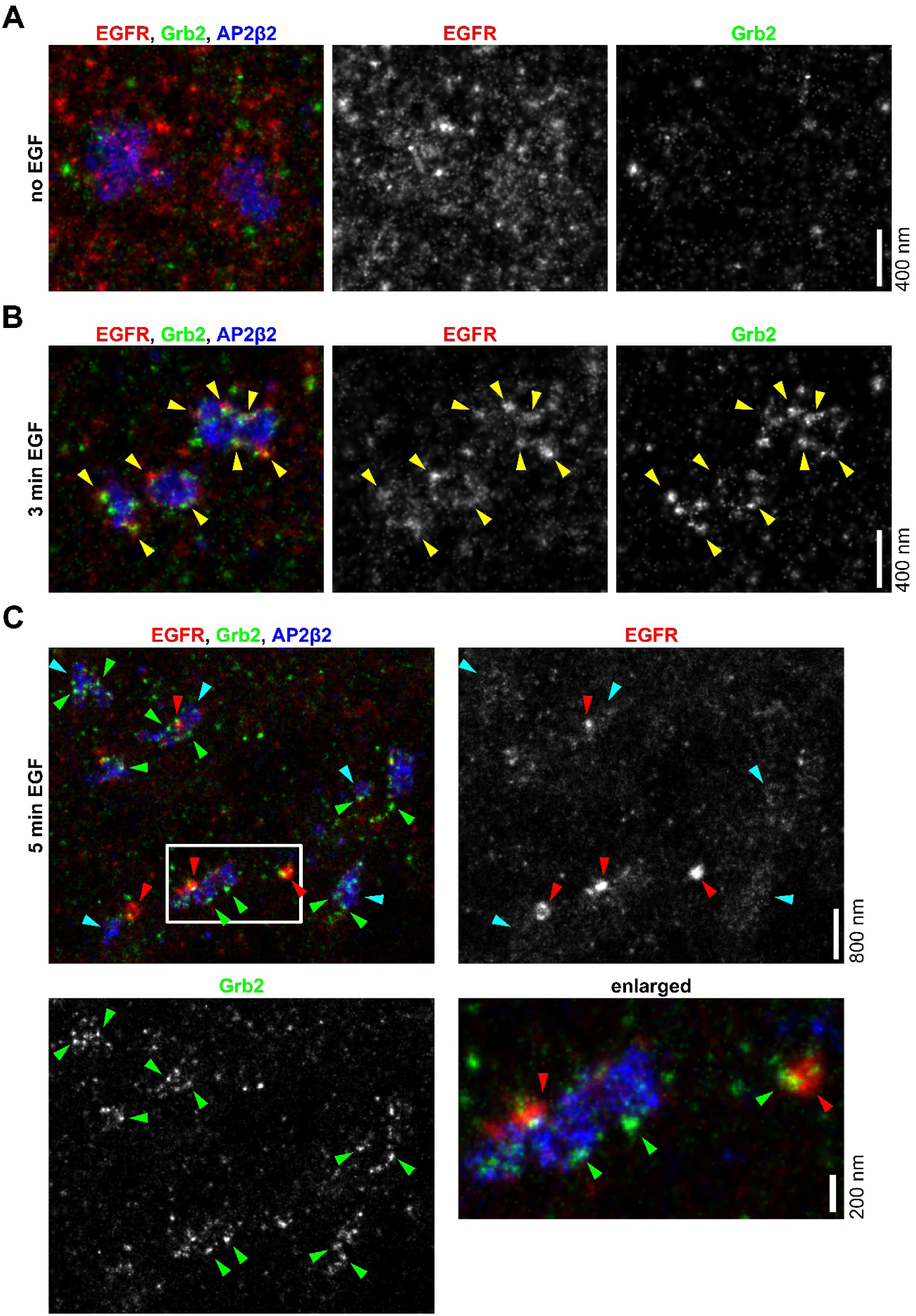
Localizations of EGFR and Grb2 before and after EGFR stimulation at 37℃. (**A**) Before the stimulation, EGFR was spread over the plasma membrane. Grb2 was not localized in the CCSs. (**B**) After 3 min of the stimulation, EGFR and Grb2 were colocalized in clusters at the CCS rims (yellow arrowheads). (**C**) After 5 min of the stimulation, EGFR clusters decreased at CCS rims (cyan arrowheads). EGFR accumulated in a round shape about 200 nm in size (red arrowheads), where Grb2 were partially localized. Many Grb2 clusters remained at the CCS rim (green arrowheads). These IRIS images were reconstructed by Gaussian rendering with mean localizations at 5 nm sized pixel.

**Extended Data Fig. 5.**
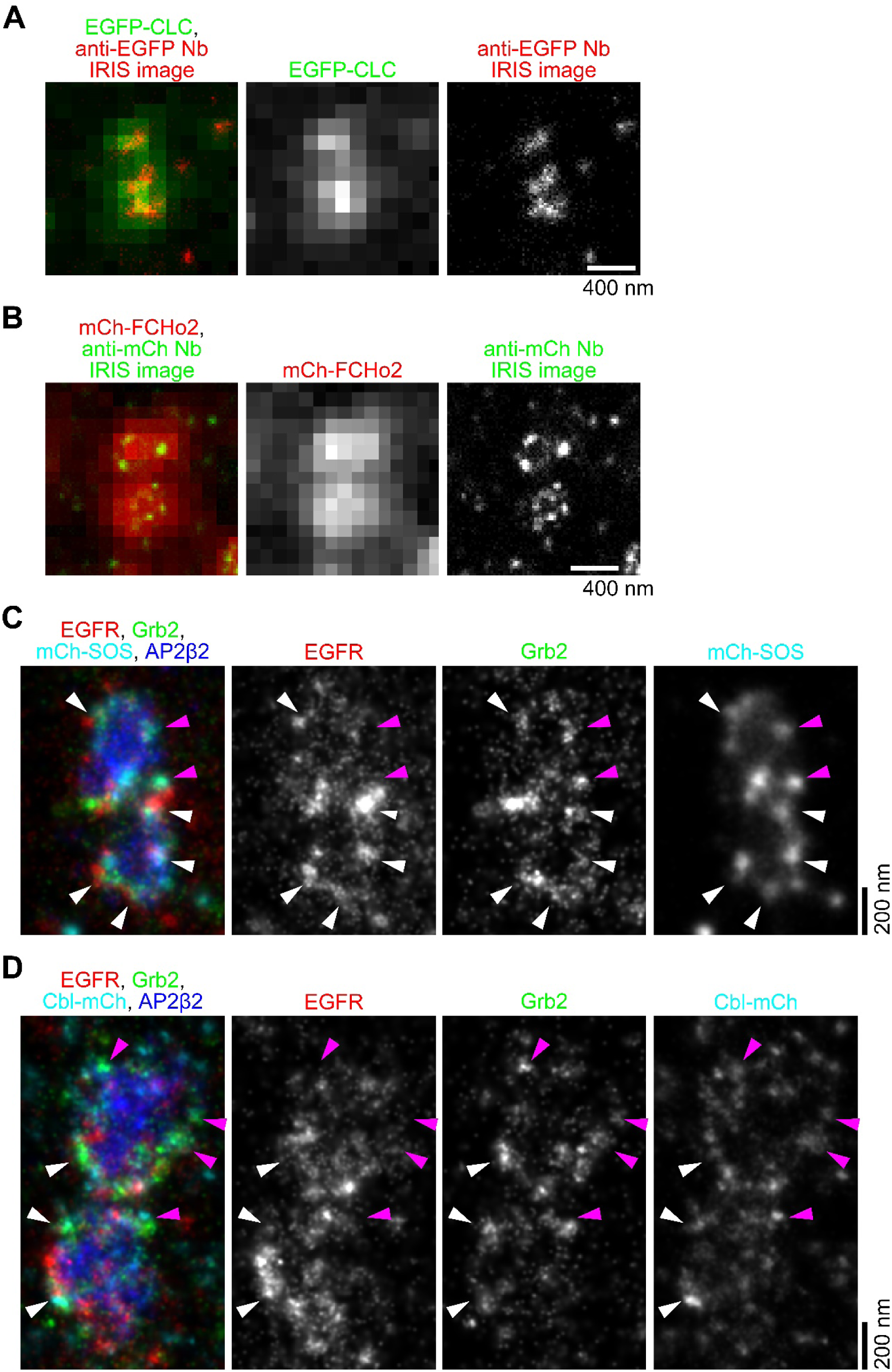
Localizations of EGFR, Grb2 and SOS1 or cCbl at CCS rims. (**A**) The fluorescence image of EGFP-CLC and IRIS image using anti-EGFP nanobody (Nb)-tagRFP. (**B**) The fluorescence image of mCh-FCHo2 and IRIS image using anti-mCh Nb-EGFP. These IRIS images were reconstructed by integrating localizations at 20 nm sized pixel. (**C**, **D**) Colocalization of mCh-SOS1 (**D**) and cCbl-mCh (**D**) with EGFR and Grb2 clusters at CCS rims. HeLa cells expressing mCh-SOS1 or cCbl-mCh were stimulated with 100 ng/ml EGF for 3 min at 37℃. These IRIS imaging were performed with anti-mCh Nb-EGFP, anti-EGFR Fab probe, anti-Grb2 Fab probe and anti-AP2β2 Fab probe. In these IRIS images, the colocalizations of EGFR and Grb2 with SOS1 or cCbl (white arrowheads) and the colocalizations of Grb2 with SOS1 or cCbl (magenta arrowheads) were observed. These IRIS images were reconstructed by Gaussian rendering with mean localizations at 5 nm sized pixel.

**Extended Data Fig. 6.**
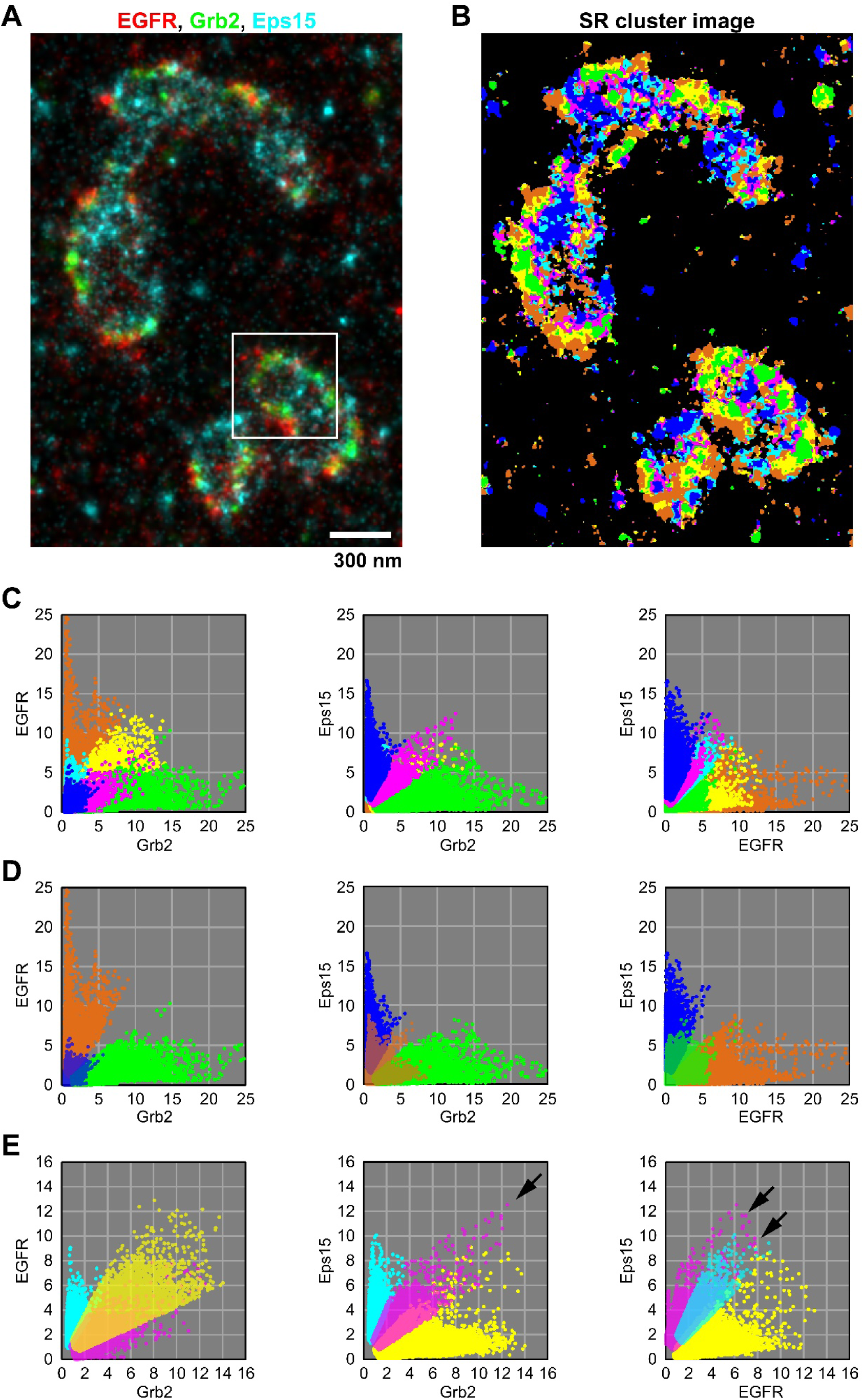
Merged IRIS image and SR cluster image of CCSs around the CCS shown in Fig. 4A. (**A**, **B**) Merged IRIS image of EGFR (red), Grb2 (green) and Eps15 (cyan) (**A**) and their SR cluster image (**B**). The colors of the pixels in the SR cluster image are the same as the colors used in Fig. 4A. The CCS in the boxed area is shown in Fig. 4A. The IRIS image was reconstructed by Gaussian rendering with mean localizations at 5 nm sized pixel. (**C**) Scatter plots of z-score intensities of the targets at the pixels in the merged IRIS image of EGFR, Grb2 and Eps15. The mean z-score intensities of the targets in the colored pixels and the correlation coefficients between the targets are shown in Fig. 4A. The z-score intensities were plotted between Grb2 and EGFR (left) or Eps15 (center), and between EGFR and Eps15 (right). (**D**) Scatter plots for comparing EGFR-dominant (orange), Grb2-dominant (green) and Eps15-dominant (blue) pixels. (**E**) Scatter plots for comparing Grb2-EGFR dominant (yellow), Grb2- Eps15 dominant (magenta) and EGFR-Eps15 dominant (cyan) pixels. The scatter plots are the same graphs shown in **Fig. 4D–F**.

**Extended Data Fig. 7.**
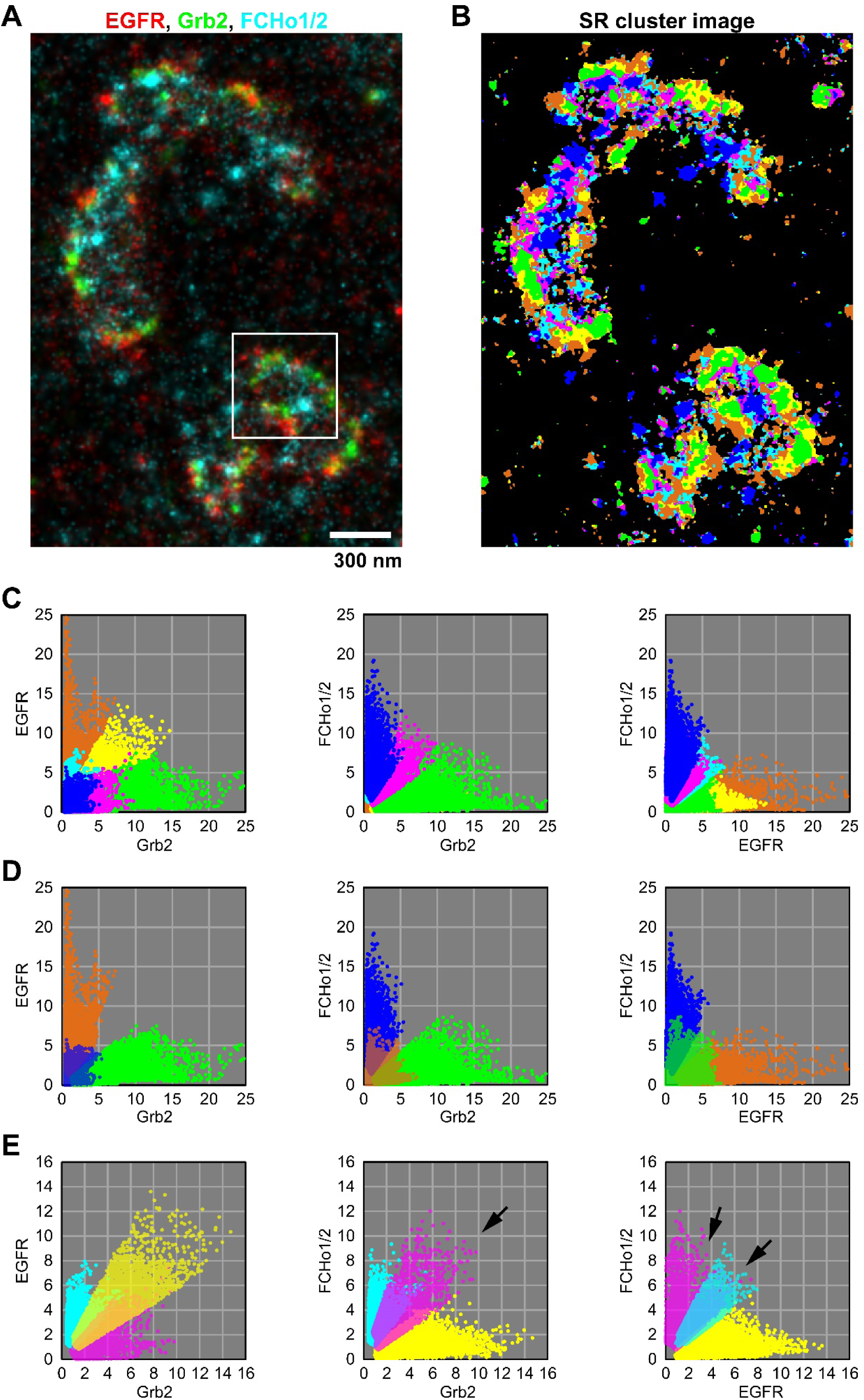
Merged IRIS image and SR cluster image of CCSs around the CCS shown in Fig. 4B. (**A**, **B**) Merged IRIS image of EGFR (red), Grb2 (green) and FCHo1/2 (cyan) (**A**) and their SR cluster image (**B**). The colors of the pixels in the SR cluster image are the same as the colors used in Fig. 4B. The CCS in the boxed area is shown in Fig. 4B. The IRIS image was reconstructed by Gaussian rendering with mean localizations at 5 nm sized pixel. (**C**) Scatter plots of z-score intensities of the targets at the pixels in the merged IRIS image of EGFR, Grb2 and FCHo1/2. The mean z-score intensities of the targets in the colored pixels and the correlation coefficients between the targets are shown in Fig. 4B. The z-score intensities were plotted between Grb2 and EGFR (left) or FCHo1/2 (center), and between EGFR and FCHo1/2 (right). (**D**) Scatter plots for comparing EGFR-dominant (orange), Grb2-dominant (green) and FCHo1/2- dominant (blue) pixels. (**E**) Scatter plots for comparing Grb2-EGFR dominant (yellow), Grb2-FCHo1/2 dominant (magenta) and EGFR-FCHo1/2 dominant (cyan) pixels.

**Extended Data Fig. 8.**
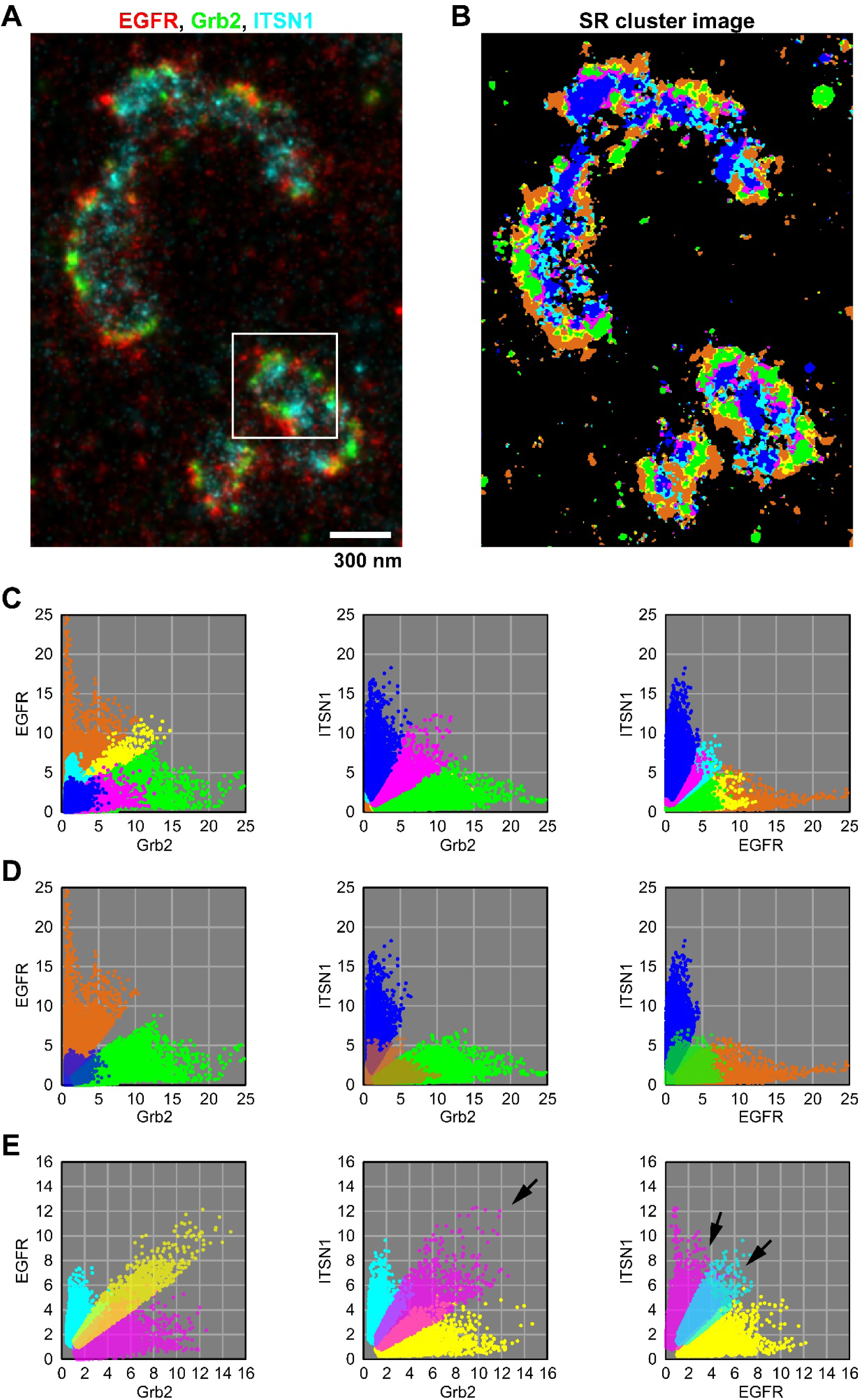
Merged IRIS image and SR cluster image of CCSs around the CCS shown in Fig. 4C. (**A**, **B**) Merged IRIS image of EGFR (red), Grb2 (green) and ITSN1 (cyan) (**A**) and their SR cluster image (**B**). The colors of the pixels in the SR cluster image are the same as the colors used in Fig. 4C. The CCS in the boxed area is shown in Fig. 4C. The IRIS image was reconstructed by Gaussian rendering with mean localizations at 5 nm sized pixel. (**C**) Scatter plots of z-score intensities of the targets at the pixels in the merged IRIS image of EGFR, Grb2 and ITSN1. The mean z-score intensities of the targets in the colored pixels and the correlation coefficients between the targets are shown in Fig. 4C. The z-score intensities were plotted between Grb2 and EGFR (left) or ITSN1 (center), and between EGFR and ITSN1 (right). (**D**) Scatter plots for comparing EGFR-dominant (orange), Grb2-dominant (green) and ITSN1-dominant (blue) pixels. (**E**) Scatter plots for comparing Grb2-EGFR dominant (yellow), Grb2- ITSN1 dominant (magenta) and EGFR-ITSN1 dominant (cyan) pixels.

**Extended Data Fig. 9.**
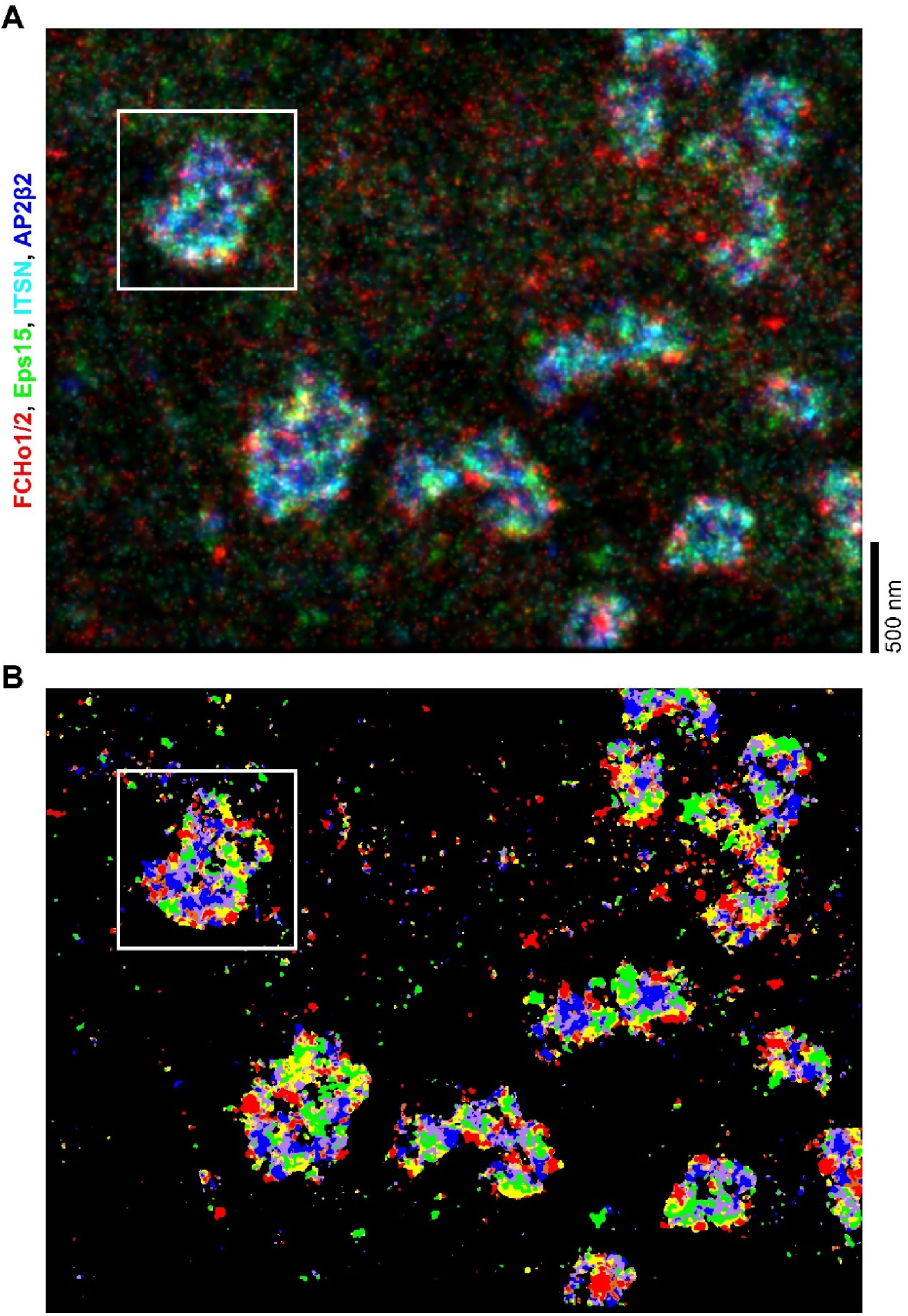
Merged IRIS image of FCHo1/2 (red), Eps15 (green), ITSN1 (cyan) and AP2β2 (blue) (A) and the SR cluster image (B). The colors of the pixels in the SR cluster image are the same as the colors used in Fig. 5B. The CCS in the boxed area is shown in Fig. 5. The IRIS image was reconstructed by Gaussian rendering with mean localizations at 5 nm sized pixel.

**Extended Data Fig. 10.**
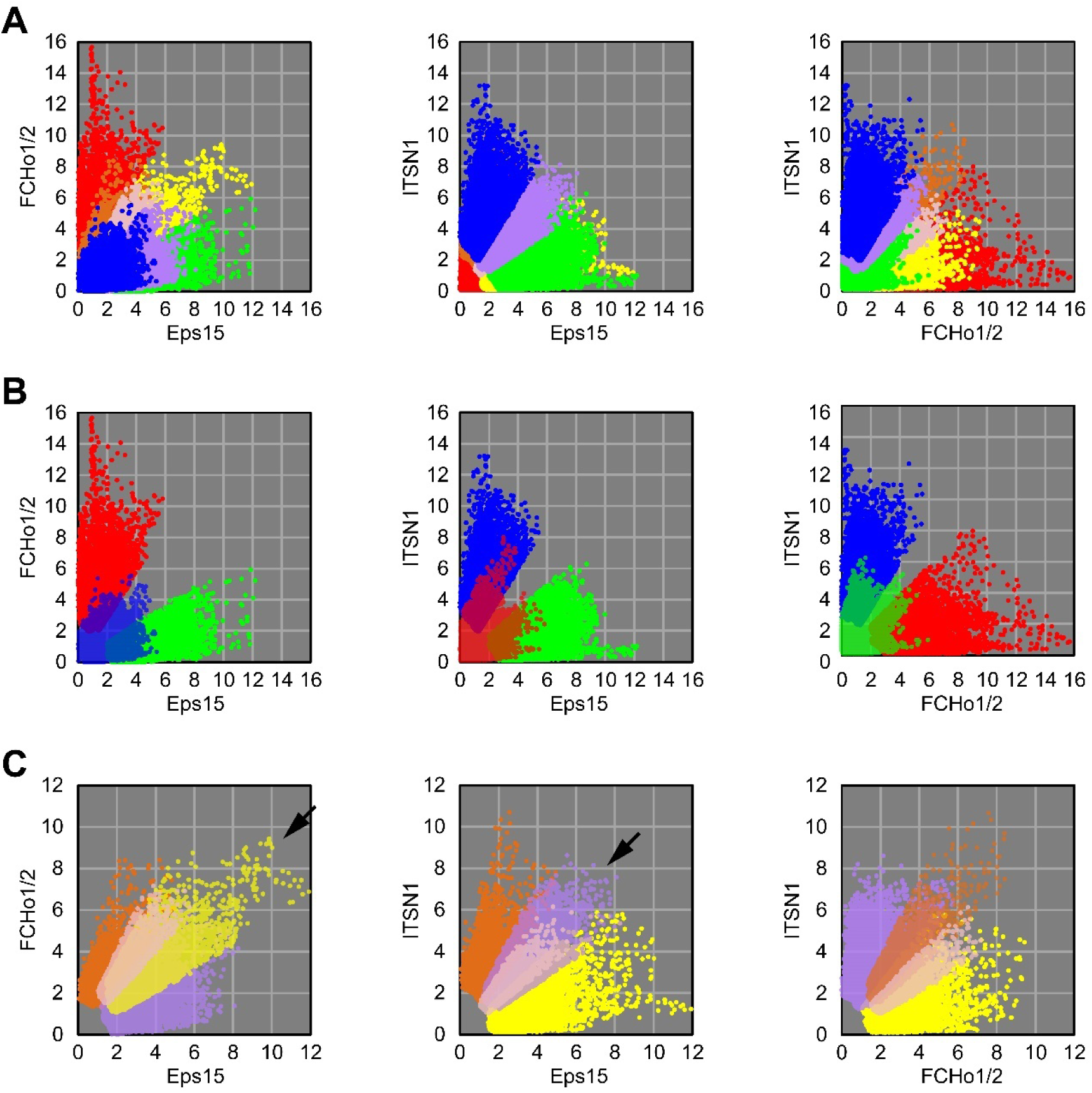
Correlation analysis of the colored pixels shown in Fig. 5 and Extended Data Fig. 9. (**A**) Scatter plots of z-score intensities of the targets at the pixels in the merged IRIS image of FCHo1/2, Eps15 and ITSN1. The z-score intensities were plotted between Eps15 and FCHo1/2 (left) or ITSN1 (center), between FCHo1/2 and ITSN1 (right). (**B**) Scatter plots for comparing FCHo1/2-dominant (red), Eps15-dominant (green) and ITSN1-dominant (blue) pixels (**C**) Scatter plots for comparing FCHo1/2- Eps15 dominant (yellow), FCHo1/2-ITSN1 dominant (orange), FCHo1/2-Eps15-ITSN1 dominant (pale pink) and Eps15-ITSN1 dominant (purple) pixels.

**Supplementary Fig. 1.**
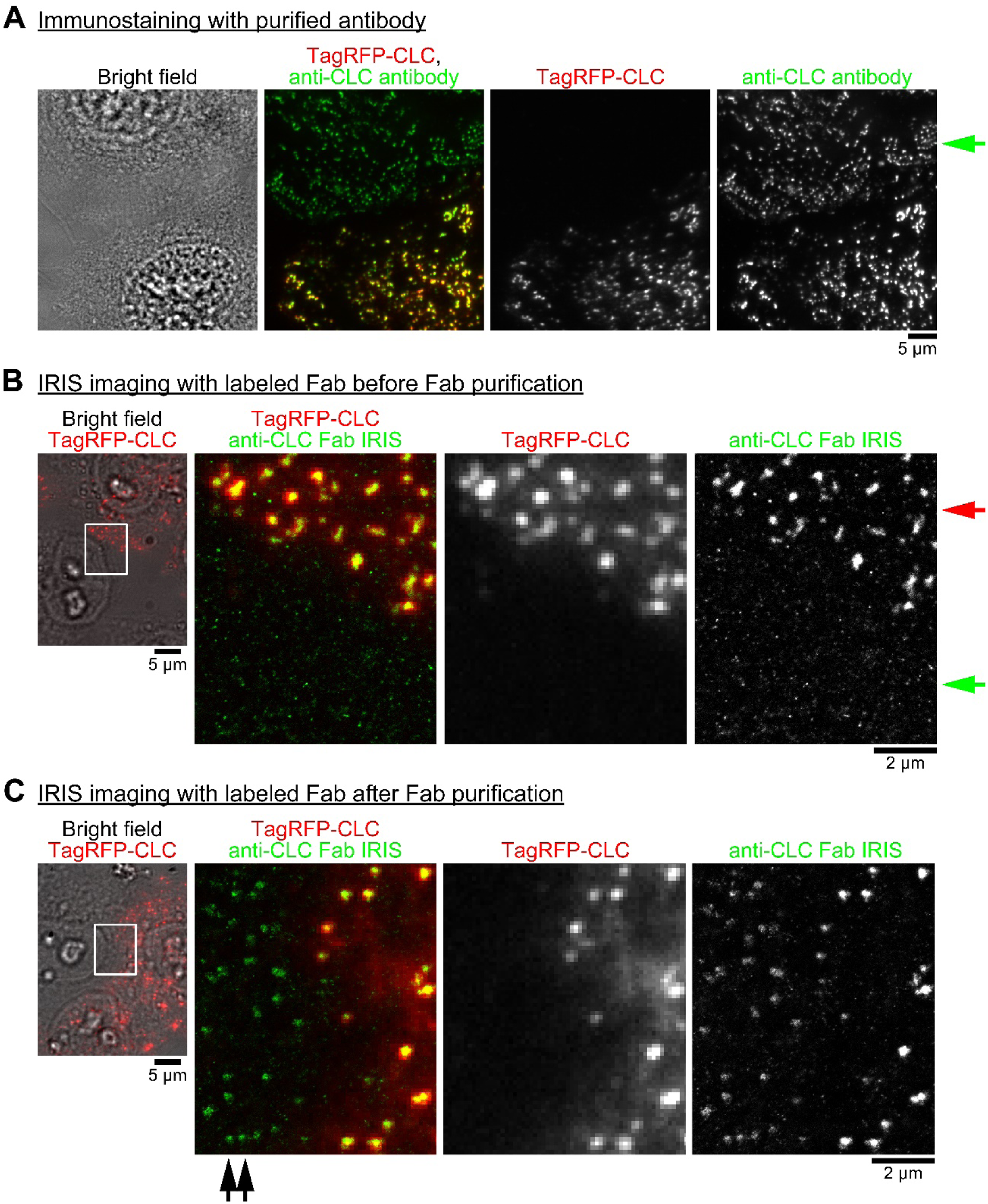
Development of a method for producing Fab probe from antiserum using anti-clathrin light chain (CLC) antiserum (see Methods). (A) Immunostaining with anti-CLC antibodies purified by affinity chromatography. The merged image of TagRFP-CLC (red) and anti-CLC antibodies (green). HeLa cells were stained with the antibodies to find the eluted fraction capable of staining endogenous CLC. Endogenous CLC was localized in spots in non-expressing cell similar to expressed CLC (green arrow). (B) IRIS imaging of CLC using fluorescent dye (Dylight 488)-conjugated Fab probe. The merged image of the bright field image (gray) and the fluorescent image of TagRFP-CLC (red) (leftmost). The merged image of the fluorescent image of TagRFP- CLC (red) and the IRIS image using the Fab probe (green) in the boxed area in the leftmost image (three images on the right). The purified anti-CLC antibodies were labeled while bound to CLC-conjugated resin to protect its antigen-binding site, which stained endogenous CLC with less non-specific signals by immunostaining. Their Fab fragments were generated by papain digestion. In the IRIS image using the Fab probe, CLC was visualized in the CLC-expressing cell (red arrow). But endogenous CLC was not visualized in the non-expressing cell, where there were non-specific signals (green arrow). (C) IRIS imaging of CLC using the Fab probe after Fab purification. The merged image of the bright field image (gray) and the fluorescent image of TagRFP-CLC (red) (leftmost). The merged image of the fluorescent image of TagRFP-CLC (red) and the IRIS image using the purified Fab probe (green) in the boxed area in the leftmost image (three images on the right). To reduce the non-specific signals of the Fab probe, the Fab probe was further purified with CLC-conjugated resin. The IRIS image using the purified Fab probe showed endogenous CLC in the non-expressing cell (double arrows).

**Supplementary Fig. 2.**
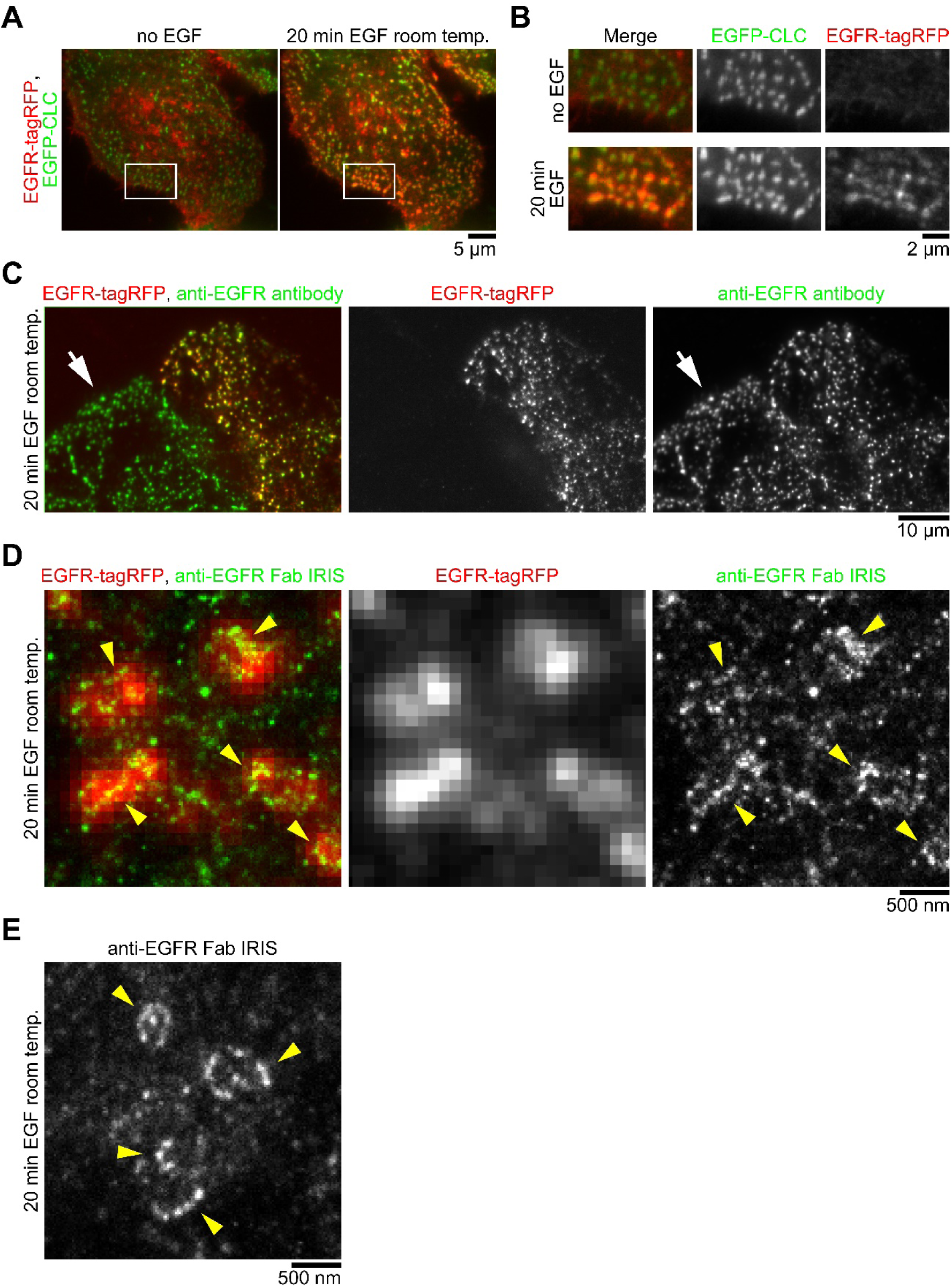
Preparation of anti-EGFR Fab probe. (**A**) Time-lapse imaging of EGFR-tagRFP and EGFP-CLC. HeLa cells before and after stimulation with 100 ng/ml EGF for 20 min at room temperature for the delayed endocytosis of EGFR. (**B**) Enlarged images in the boxed area shown in **A**. Even 20 min after the stimulation, EGFR was accumulating in CCSs. (**C**) Immunostaining with anti-EGFR antibodies purified by affinity chromatography. The merged image of EGFR-tagRFP (red) and anti-EGFR antibodies (green). After EGF stimulation, HeLa cells were stained with the antibodies to find the eluted fraction capable of staining endogenous EGFR. Endogenous EGFR was localized in spots in non-expressing cell similar to expressed EGFR (arrow). (**D**) IRIS imaging using anti-EGFR Fab probe in EGFR-tagRFP-expressing cells. The merged image of the fluorescent image of EGFR-tagRFP (red) and the IRIS image (green). (**E**) IRIS imaging of using anti-EGFR Fab probe in non-expressing cells. EGFR was localized in cluster along the rim of some region with a size of several hundred nanometers (presumably CCSs) (**D**, **E**, yellow arrowheads).

**Supplementary Fig. 3.**
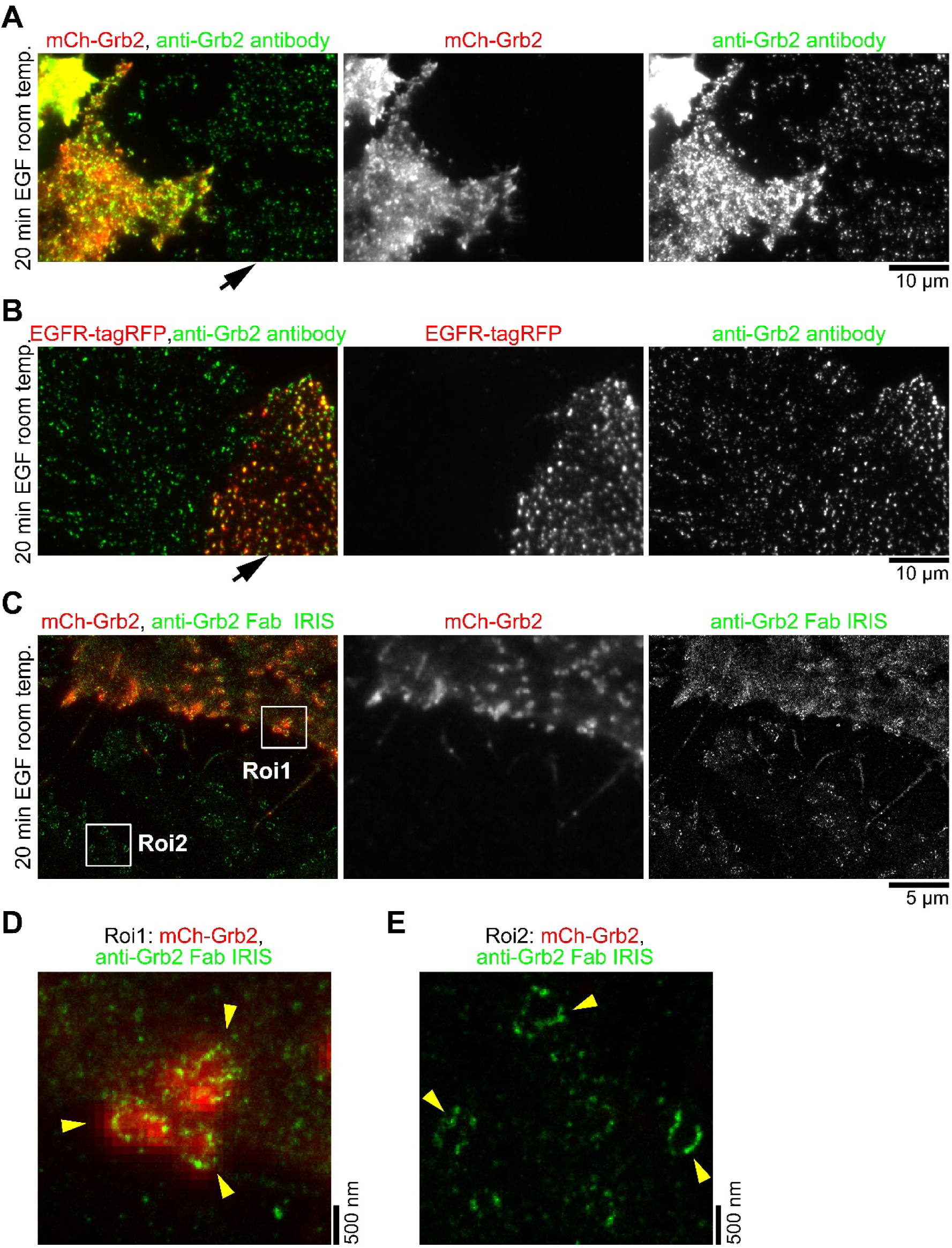
Preparation of anti-Grb2 Fab probe. (**A**, **B**) Immunostaining with anti-Grb2 antibodies purified by affinity chromatography. The merged image of anti- Grb2 antibodies (green) and mCherry (mCh)-Grb2 (red) (**A**) or EGFR-tagRFP (red) (**B**). HeLa cells were stimulated with 100 ng/ml EGF for 20 min at room temperature. The cells were stained with the antibodies to find the eluted fraction capable of staining endogenous Grb2. Endogenous Grb2 was localized in spots in non-expressing cell (**A**, arrow). The spots of Grb2 were colocalized with EGFR-tagRFP (**B**, arrow). As a supplement, this Grb2 antibody could not stain Grb2 with mCh fused to its C-terminus. (**C–E**) IRIS imaging using anti-Grb2 Fab probe. The merged image of the fluorescent image of mCh-Grb2 (red) and the IRIS image (green). Grb2 were localized along the rim of some region with a size of several hundred nanometers (presumably CCSs) in mCh- Grb2-expressing cell (**D**) and in non-expressing cell (**E**) shown in **C** (yellow arrowheads).

**Supplementary Fig. 4.**
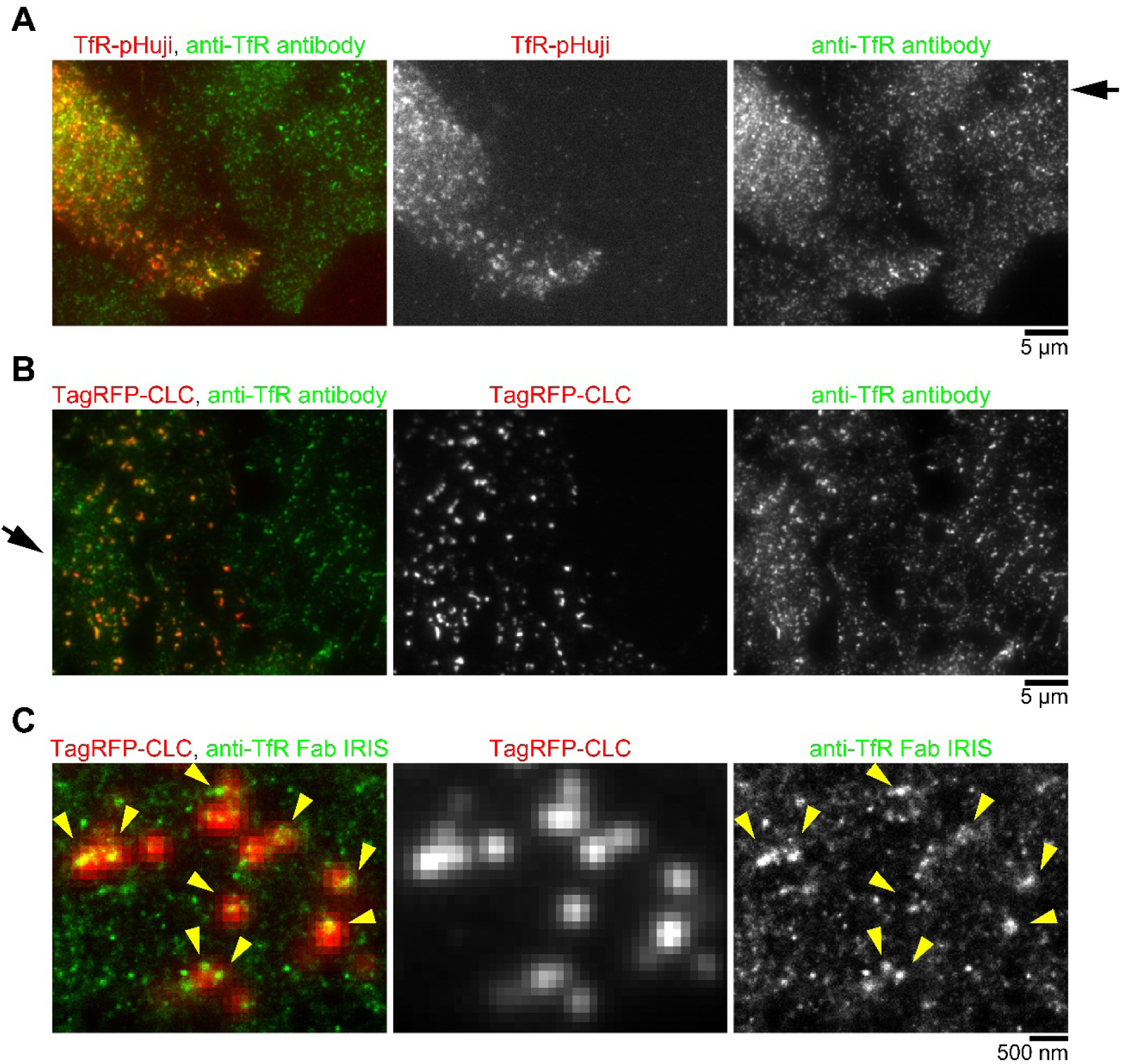
Preparation of anti-transferrin receptor (TfR) Fab probe. (**A**, **B**) Immunostaining with anti-TfR antibodies purified by affinity chromatography. The merged image of anti-TfR antibodies (green) and TfR-pHuji (red) (**A**) or TagRFP-CLC (red) (**B**). HeLa cells were stained with the antibodies to find the eluted fraction capable of staining endogenous TfR. Endogenous TfR was distributed in spots in non-expressing cell similar to expressed TfR (**A**, arrow). The spots of TfR were colocalized with TagRFP- CLC (**B**, arrow). (**C**) IRIS imaging using anti-TfR Fab probe. The merged image of the fluorescent image of TagRFP-CLC (red) and the IRIS image (green). The high intensity spots of TfR were localized in CCSs shown by TagRFP-CLC (yellow arrowheads).

**Supplementary Fig. 5.**
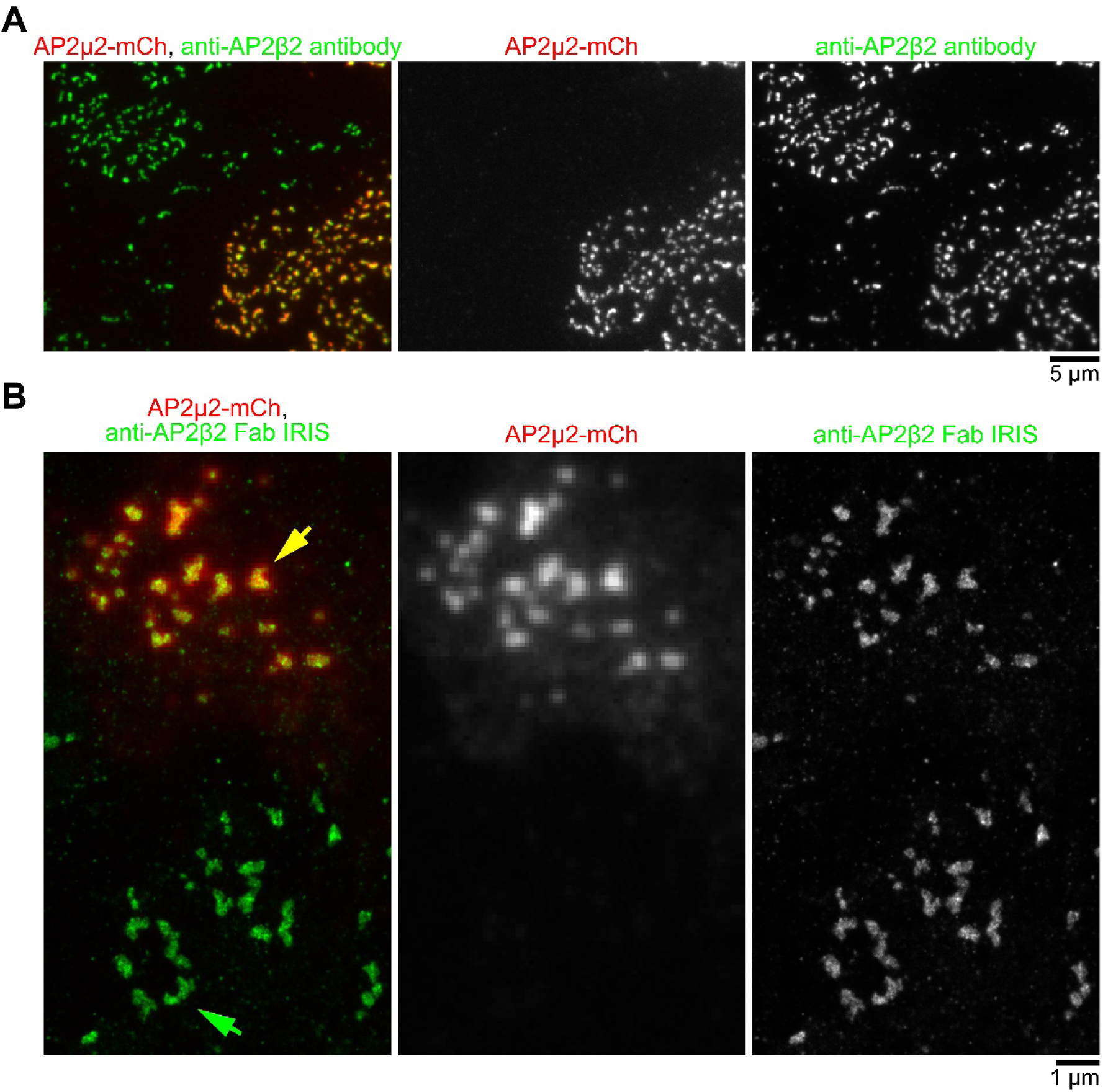
Preparation of anti-adaptor protein 2 complex beta subunit (AP2β2) Fab probe. (**A**) Immunostaining with anti-AP2β2 antibodies purified by affinity chromatography. The merged image of AP2μ2-mCh (red) and anti-AP2β2 antibodies (green). AP2β2 forms a heterotetrameric complex of α, β2, μ2 and σ2 subunits and binds to clathrin heavy chain. HeLa cells expressing AP2μ2-mCh were stained with the antibodies to find the eluted fraction capable of staining endogenous AP2β2. (**B**) IRIS imaging using anti-AP2β2 Fab probe. The merged image of the fluorescent image of AP2μ2-mCh (red) and the IRIS image (green). Endogenous AP2β2 was localized like CCSs in the non-expressing cell (green arrow) and colocalized with the accumulation of AP2μ2-mCh (yellow arrow).

**Supplementary Fig. 6.**
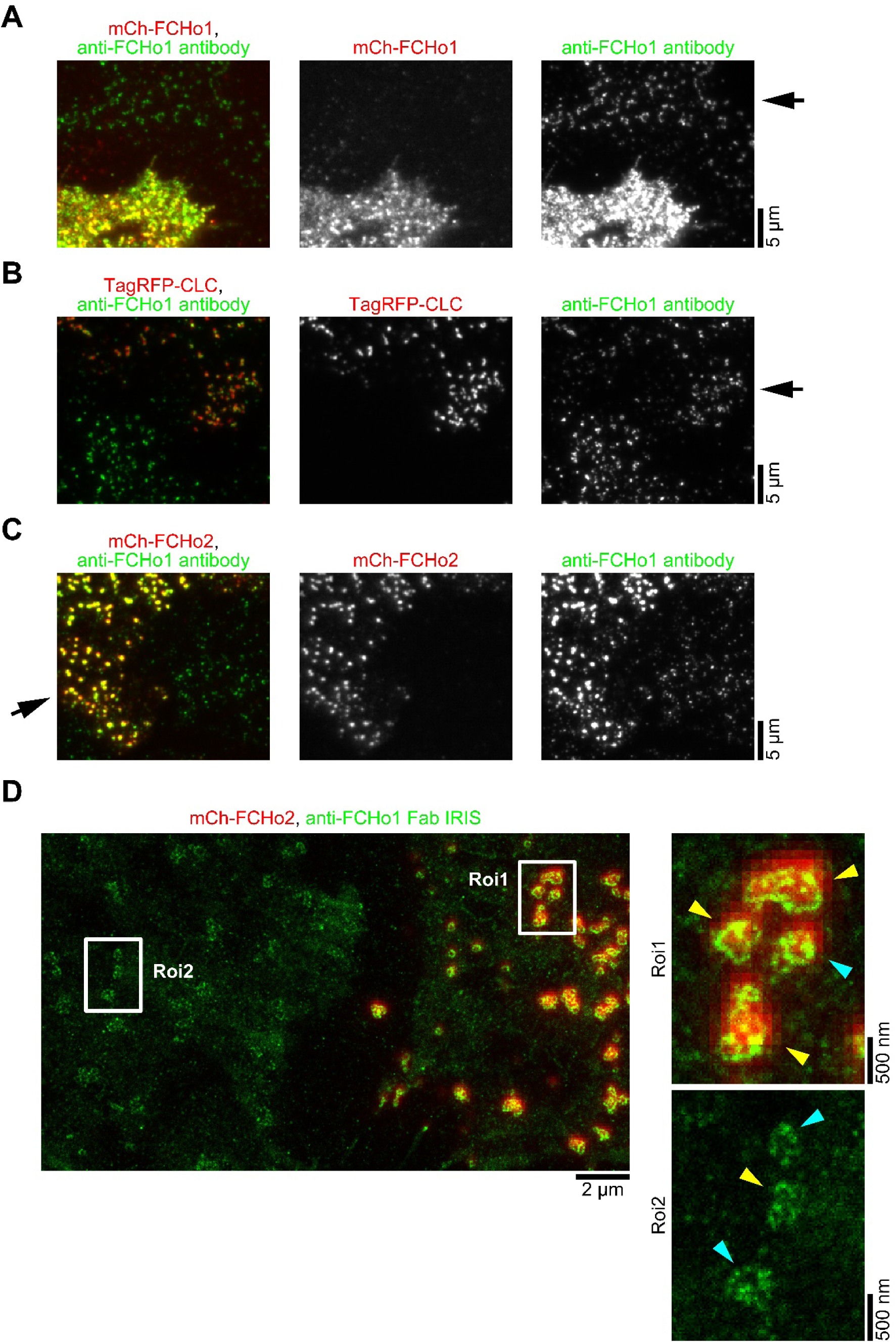
Preparation of anti-FCHo1 Fab probe. (**A–C**) Immunostaining with anti-FCHo1 antibodies purified by affinity chromatography. The merged image of anti-FCHo1 antibodies (green) and mCh-FCHo1 (red) (**A**), TagRFP-CLC (red) (**B**) or mCh-FCHo2 (red) (**C**). HeLa cells were stained with the antibodies to find the eluted fraction capable of staining endogenous FCHo1. Endogenous FCHo1 was distributed in spots in non-expressing cell similar to expressed FCHo1 (**A**, arrow). The spots of FCHo1 were colocalized with TagRFP-CLC (**B**, arrow). In addition, this anti-FCHo1 antibody stained mCh-FCHo2 (**C**, arrow), which indicates that this antibody can stain both FCHo1 and 2 (FCHo1/2). (**D**) IRIS imaging using anti-FCHo1 Fab probe. The merged image of the fluorescent image of mCh-FCHo2 (red) and the IRIS image (green). FCHo1/2 was localized along the rim of some region (presumably CCSs) with a size of several hundred nanometers in mCh-FCHo2-expressing cell (right upper image) and in non-expressing cell (right lower image) (cyan arrowheads). In addition, the multi-compartmental localizations were also observed (yellow arrowheads), which appears to be the rims of several clathrin plaques in close proximity observed by electron microscopy^12^.

**Supplementary Fig. 7.**
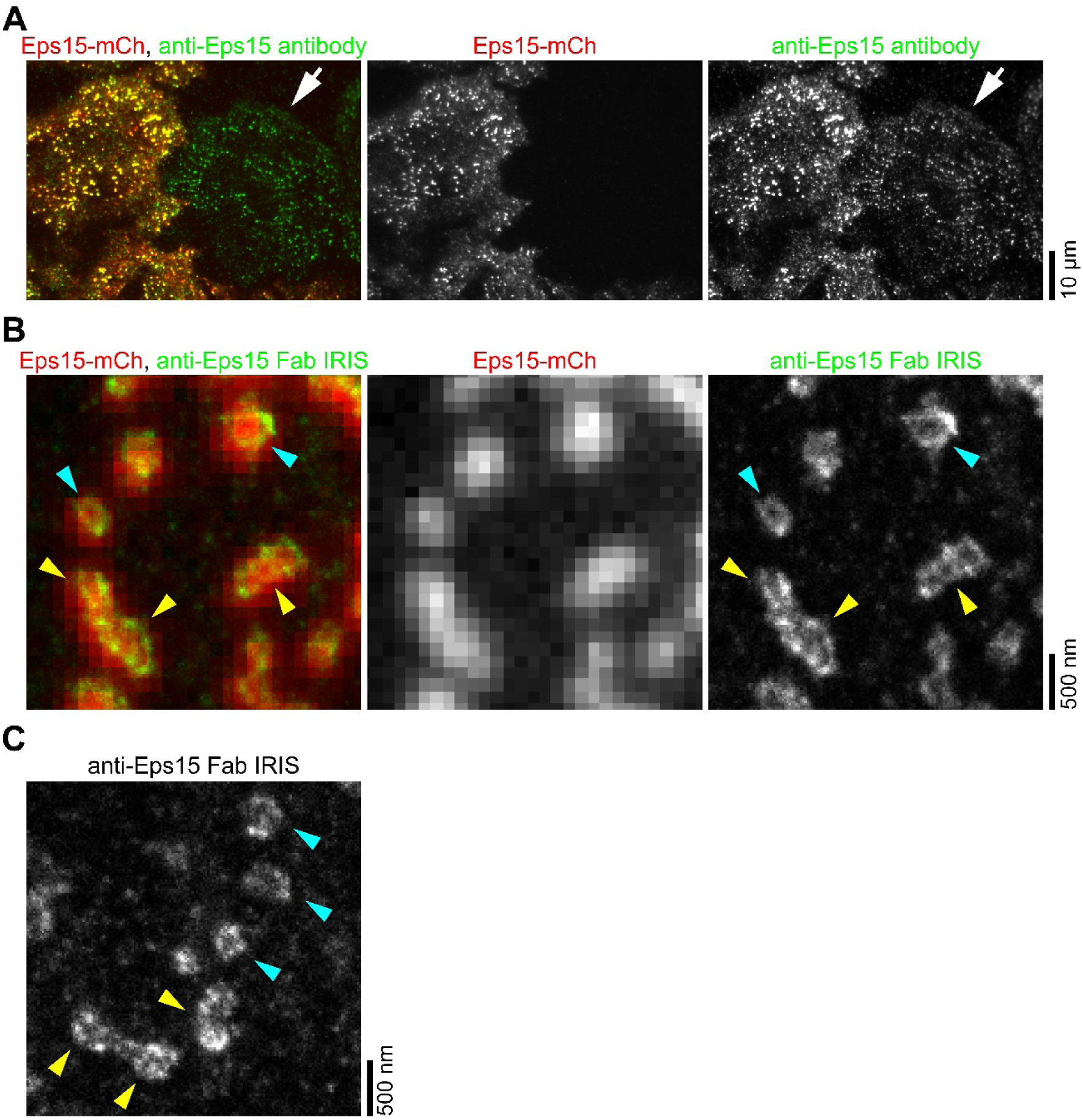
Preparation of anti-Eps15 Fab probe. (**A**) Immunostaining with anti-Eps15 antibodies purified by affinity chromatography. The merged image of Eps15-mCh (red) and anti-Eps15 antibodies (green). HeLa cells were stained with the antibodies to find the eluted fraction capable of staining endogenous Eps15. Endogenous Eps15 was distributed in spots in non-expressing cell similar to expressed Eps15 (arrow). (**B**, **C**) IRIS imaging using anti-Eps15 Fab probe. The merged image of the fluorescent image of Eps15-mCh (red) and the IRIS image (green). The expressed (**B**) and endogenous (**C**) Eps15 were visualized. These images show that Eps15 were not only localized along the rim of some region (presumably CCSs) with a size of several hundred nanometers (cyan arrowheads), but also as multi-compartmental localizations (yellow arrowheads), similar to FCHo1/2.

**Supplementary Fig. 8.**
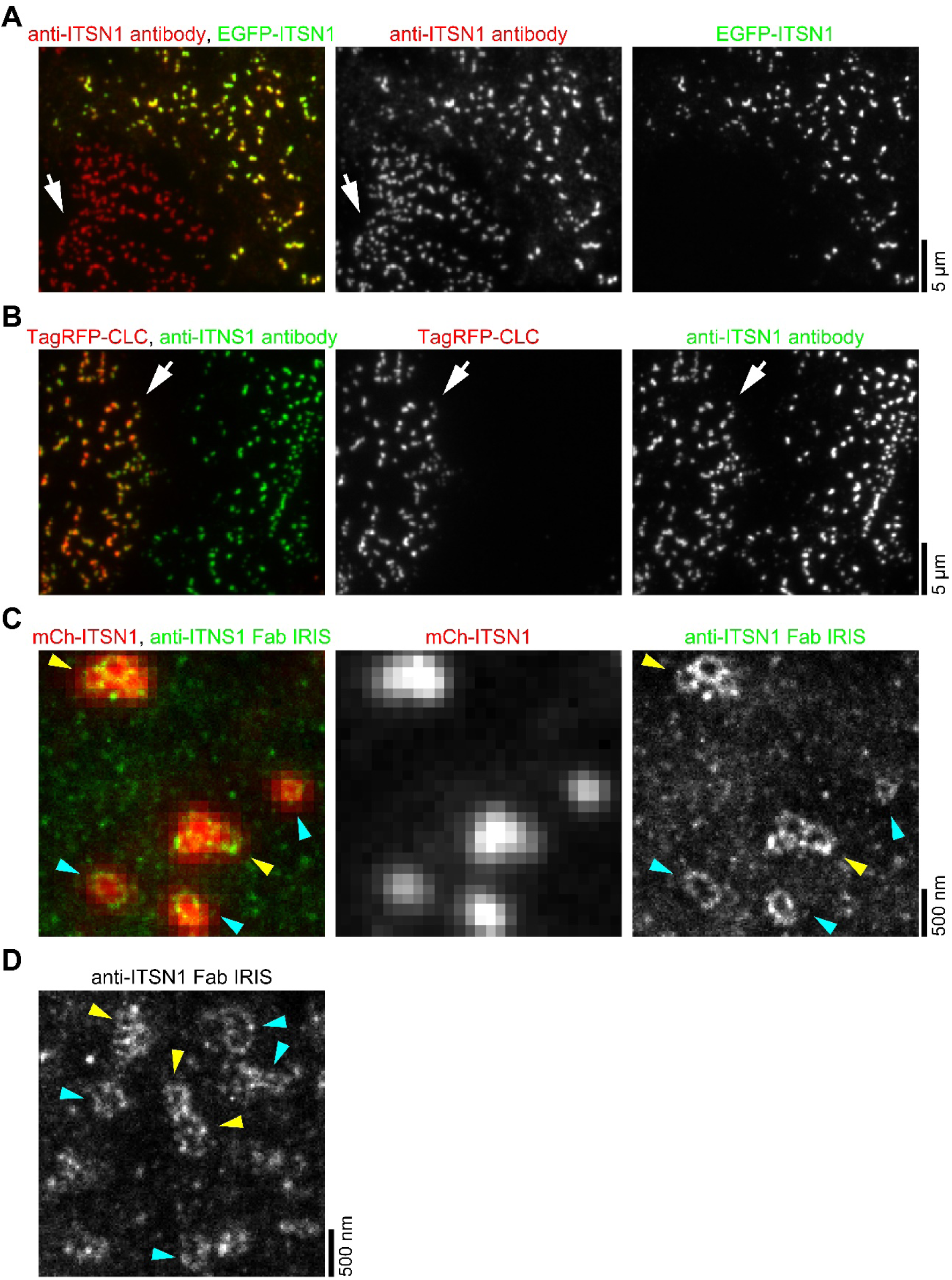
Preparation of anti-intersectin-1 (ITSN1) Fab probe. (**A**, **B**) Immunostaining with anti-ITSN1 antibodies purified by affinity chromatography. The merged image of anti-ITSN1 antibodies (red) and EGFP-ITSN1 (green) (**A**). The merged image of anti-ITSN1 antibodies (green) and TagRFP-CLC (red) (**B**). HeLa cells were stained with the antibodies to find the eluted fraction capable of staining endogenous ITSN1. Endogenous ITSN1 was distributed in spots in non-expressing cell similar to expressed ITSN1 (**A**, arrow). The spots of ITSN1 were colocalized with TagRFP-CLC (**B**, arrow). (**C**, **D**) IRIS imaging using anti-ITSN1 Fab probe. The expressed (**C**) and endogenous (**D**) ITSN1 were visualized. These images showed that ITSN1 were not only localized along the rim of some region (presumably CCSs) with a size of several hundred nanometers (cyan arrowheads), but also as multi-compartmental localizations (yellow arrowheads) , similar to FCHo1/2.

**Supplementary Fig. 9.**
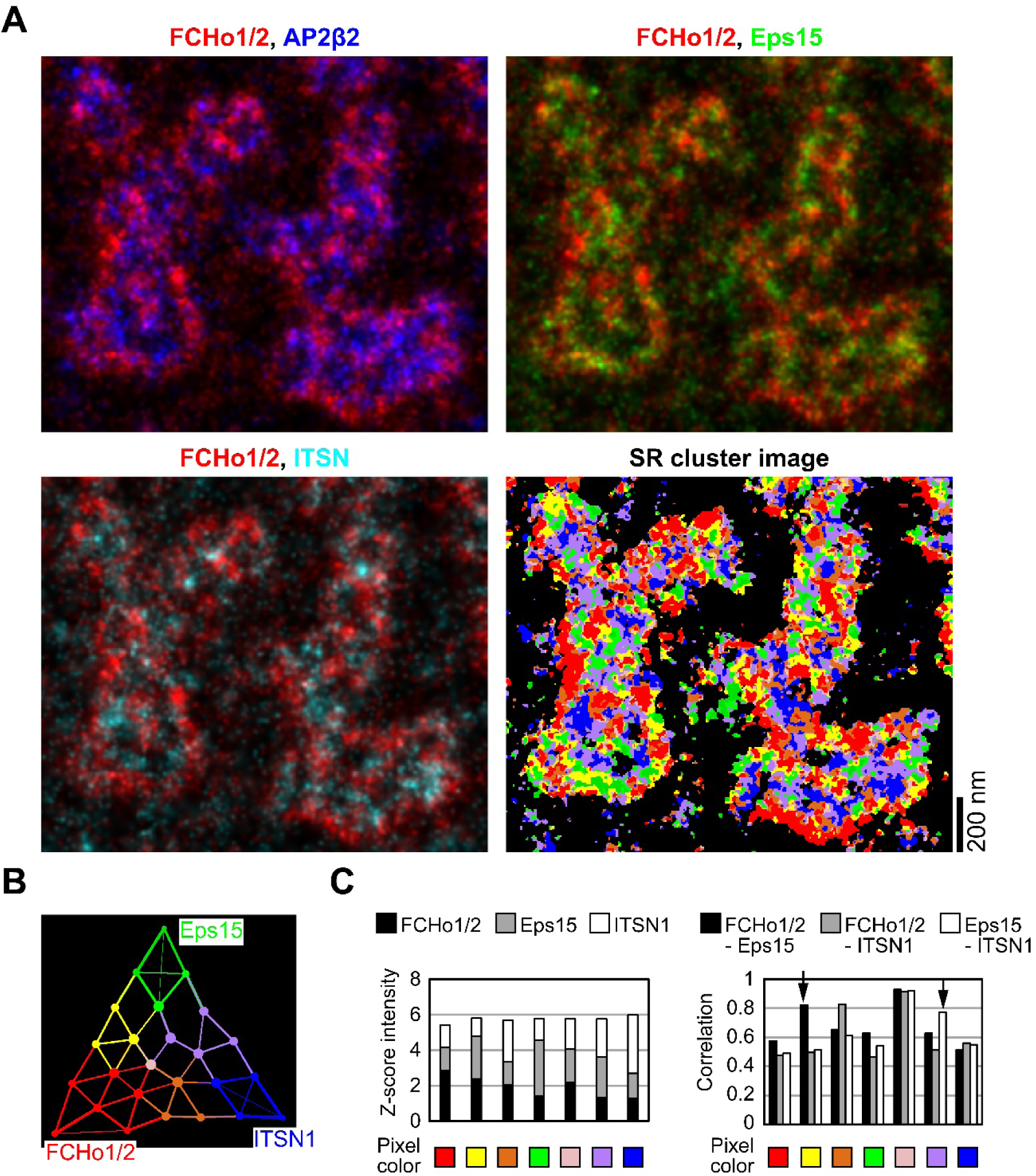
Colocalization analysis of FCHo1/2, Eps15 and ITSN1 in HeLa cells overexpressing mCh-FCHo2. (**A**) Merged IRIS images of FCHo1/2 (red), Eps15 (green), ITSN1 (cyan) and AP2β2 (blue) and the SR cluster image. The IRIS images were reconstructed by Gaussian rendering with mean localizations at 5 nm sized pixel. (**B**) PCA cluster image of FCHo1, Eps15 and ITSN1. (**C**) The mean z-score intensities of the targets in the colored pixels (left) and the correlation coefficients between the targets (right). In the cells overexpressing mCh-FCHo2, the localization of FCHo1/2 along the CCS rim was enhanced.

**Supplementary Table 1.**
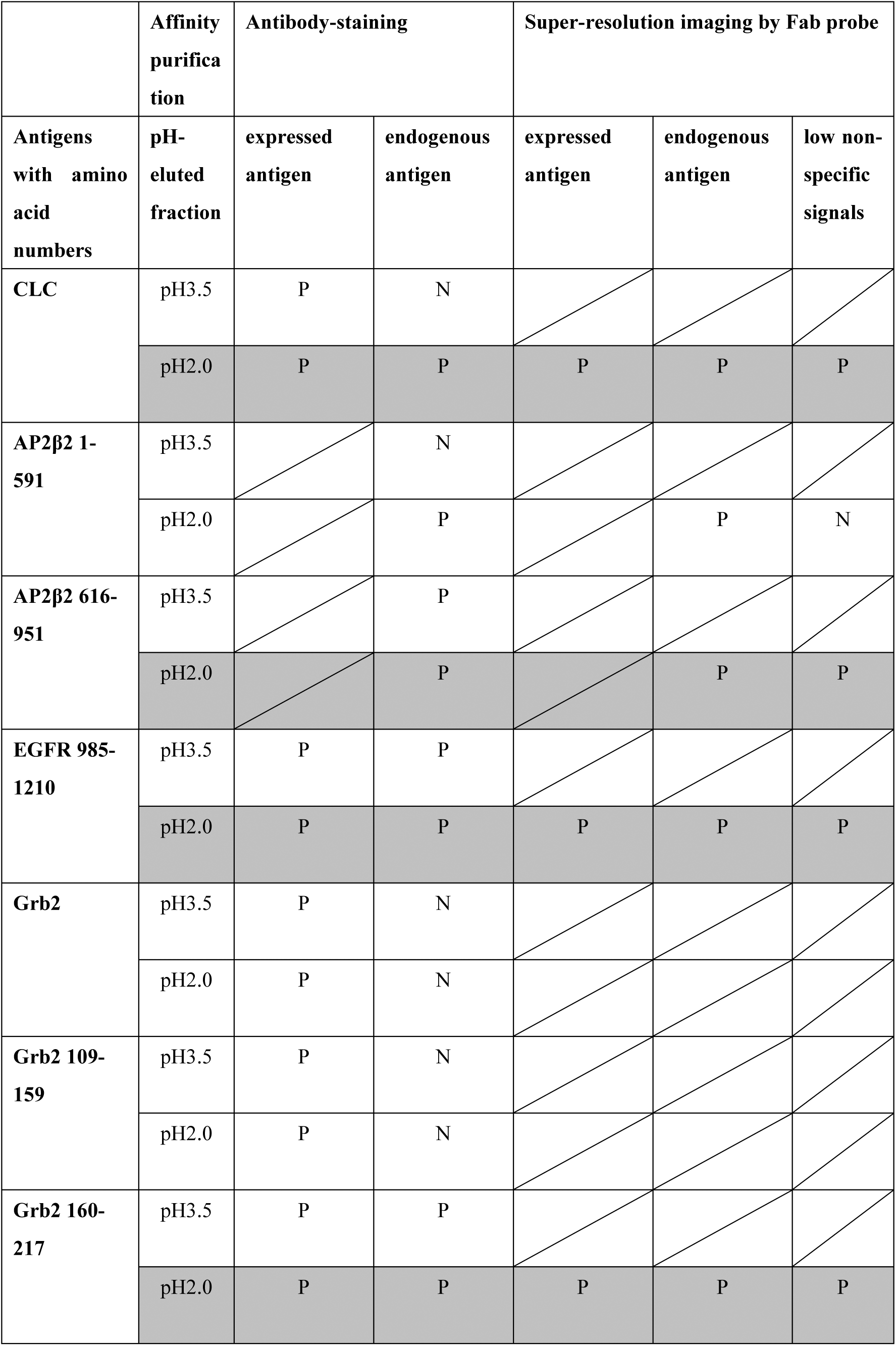
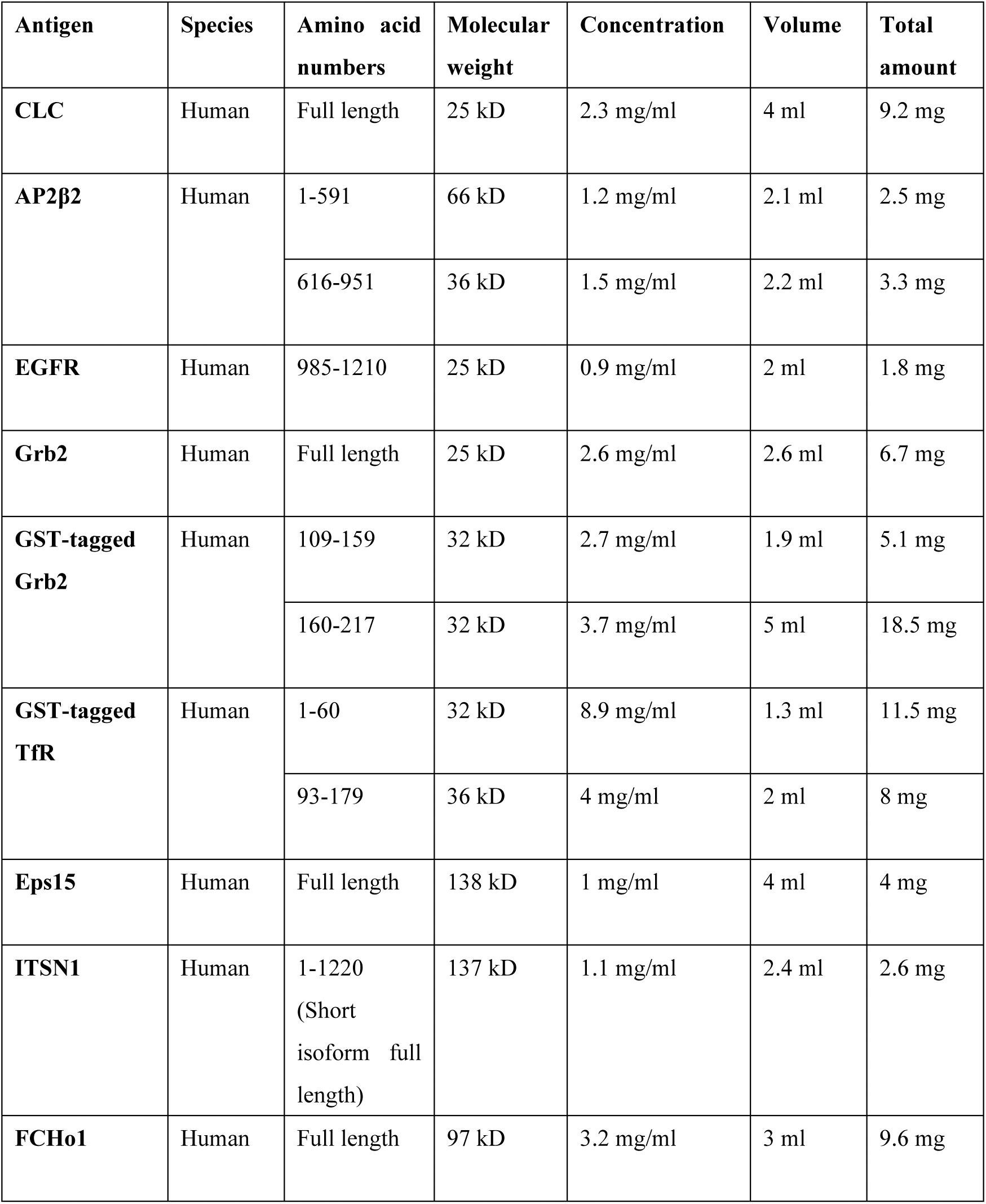
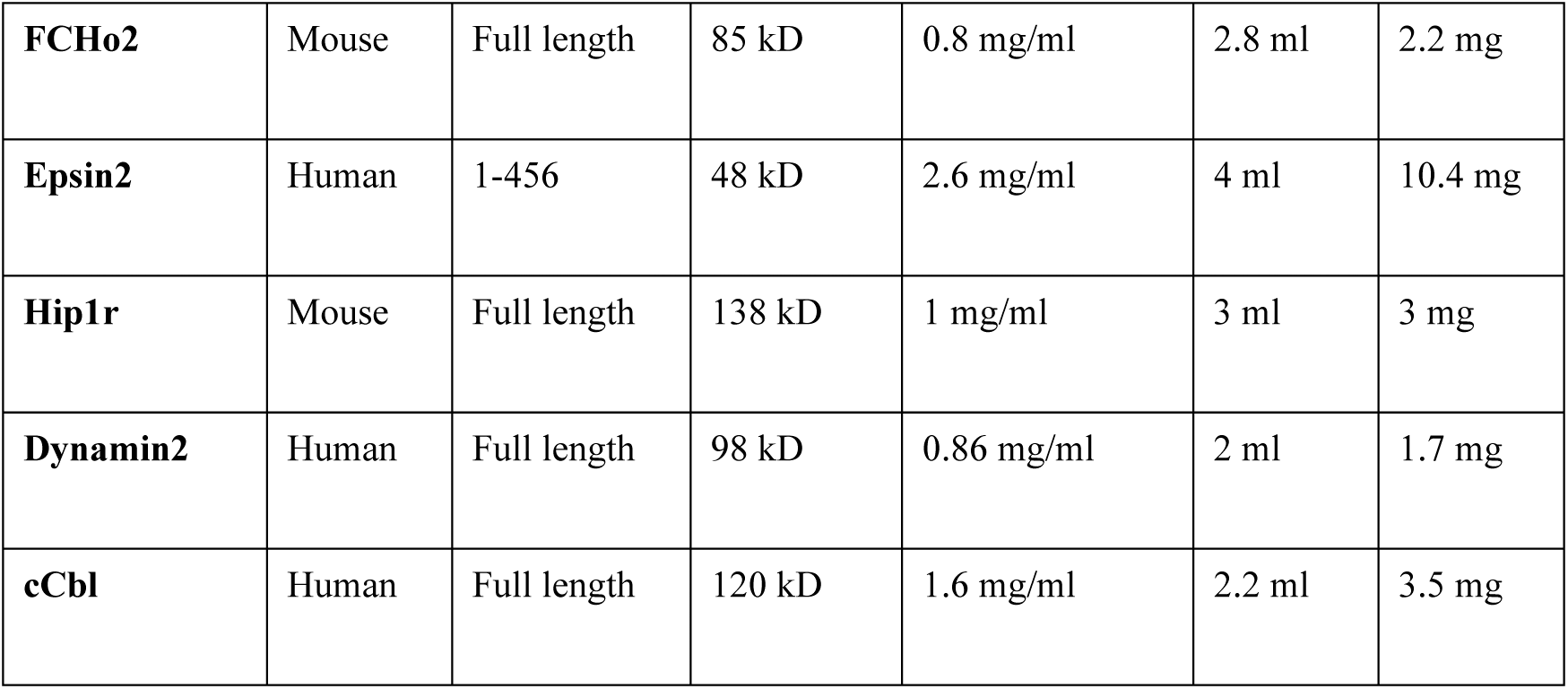
List of antibodies purified from antisera.

**Supplementary Table 2.**
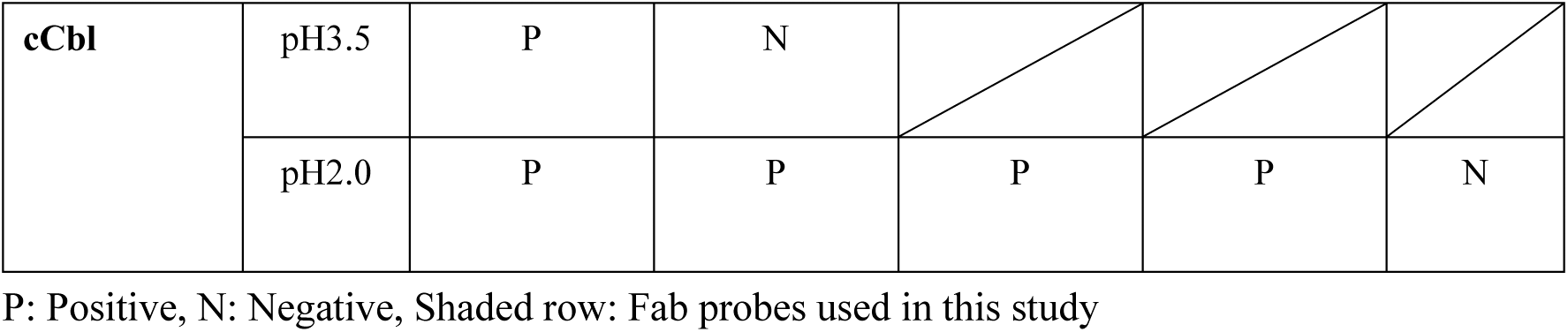
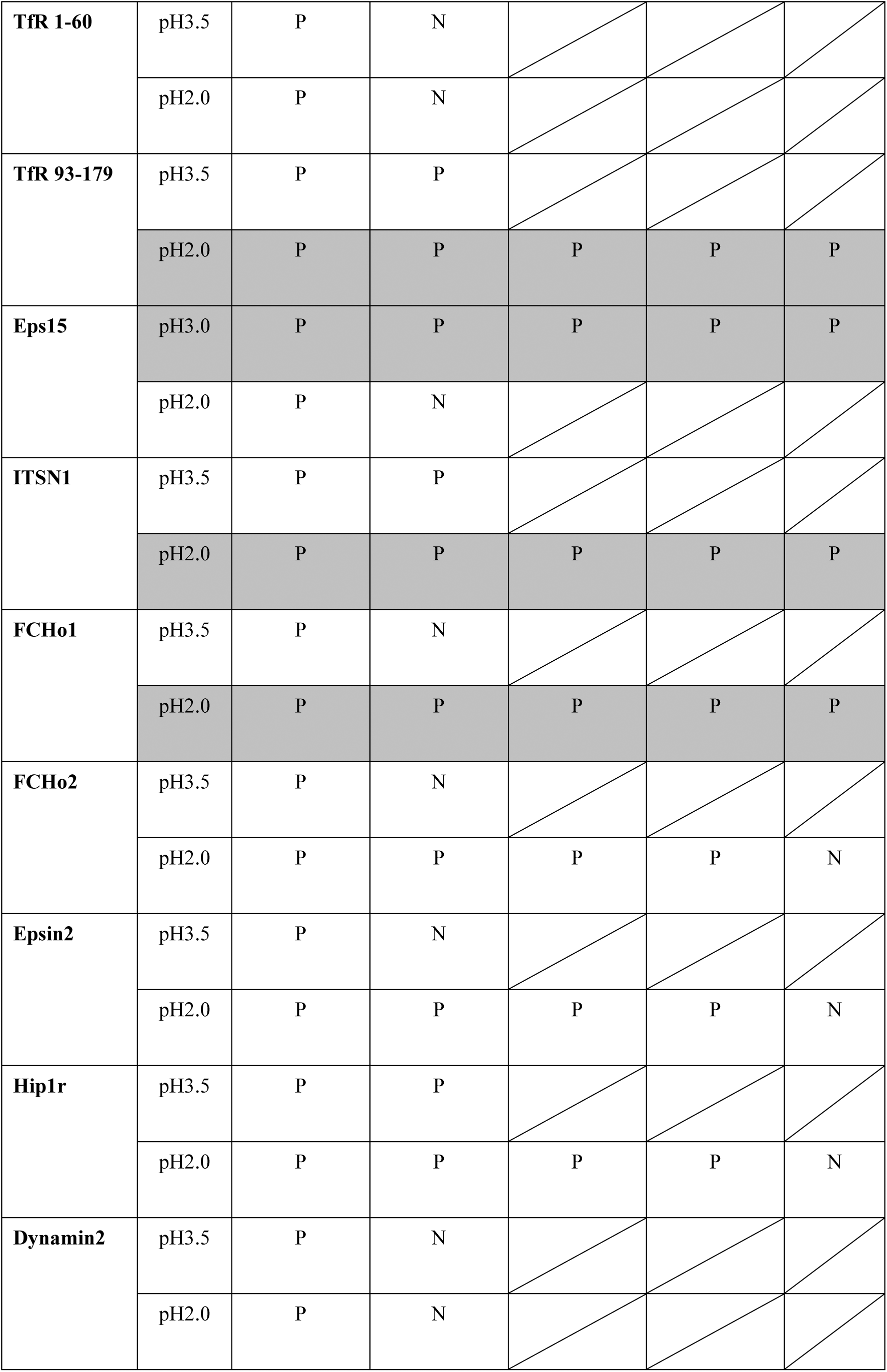
List of antigens purified from a 6L E. coli culture in one or two experiments, for FCHo2 in three or four experiments.

